# Targeting GOF p53 and c-MYC through LZK Inhibition or Degradation Suppresses Head and Neck Tumor Growth

**DOI:** 10.1101/2024.11.19.623840

**Authors:** Amy L. Funk, Meghri Katerji, Marwa Afifi, Katherine Nyswaner, Carolyn C. Woodroofe, Zoe C. Edwards, Eric Lindberg, Knickole L. Bergman, Nancy R. Gough, Maxine R. Rubin, Kamila Karpińska, Eleanor W. Trotter, Sweta Dash, Amy L. Ries, Amy James, Christina M. Robinson, Simone Difilippantonio, Baktiar O. Karim, Ting-Chia Chang, Li Chen, Xin Xu, James H. Doroshow, Ivan Ahel, Anna A. Marusiak, Rolf E. Swenson, Steven D. Cappell, John Brognard

## Abstract

The worldwide frequency of head and neck squamous cell carcinoma (HNSCC) is approximately 800,000 new cases, with 430,000 deaths annually. We determined that LZK (encoded by *MAP3K13*) is a therapeutic target in HNSCC and showed that inhibition with small molecule inhibitors decreases the viability of HNSCC cells with amplified *MAP3K13*. A drug-resistant mutant of LZK blocks decreases in cell viability due to LZK inhibition, indicating on-target activity by two separate small molecules. Inhibition of LZK catalytic activity suppressed tumor growth in HNSCC PDX models with amplified *MAP3K13*. We found that the kinase activity of LZK stabilized c-MYC and that LZK stabilized gain-of-function (GOF) p53 through a kinase-independent mechanism. Therefore, we designed proteolysis-targeting chimeras (PROTACs) and demonstrate that our lead PROTAC promotes LZK degradation and suppresses expression of GOF p53 and c-MYC leading to impaired viability of HNSCC cell lines. This research provides a strong basis for development of therapeutics targeting LZK in HNSCCs with amplification of the gene.

**One Sentence Summary:** This study establishes the kinase LZK as a therapeutic target for HNSCC through regulation of c-MYC expression.

## INTRODUCTION

Copy number alterations are frequently observed in HNSCC with 50% of patients presenting with gains of chromosome 3q (*1*) and 20% presenting with a distal amplification of chromosome 3 (3q26-3q29) (*2*). Both amplification events include *MAP3K13* encoding leucine zipper-bearing kinase (LZK), a serine-threonine protein kinase. Even with the data regarding the HNSCC genetic landscape, treatment options for HNSCC patients remain limited and include surgery, radiotherapy, platinum-based chemotherapy, and pembrolizumab (an antibody targeting programmed death-1) (*3–6*). The only approved targeted therapy for HNSCC is cetuximab, a monoclonal antibody targeting EGFR, which can increase the efficacy of chemotherapy or radiotherapy in patients (*7–9*). However, only a subset (13%) of HNSCC patients respond to cetuximab (*10*), highlighting the urgent need for new precision-based therapies for this cancer (*3, 7*).

The amplified region of chromosome 3q contains several genes known to promote cancer progression, including the transcription factor-encoding genes *SOX2* and *TP63*, and the kinase-encoding genes *TNIK* and *PIK3CA*. Of these, only PIK3CA and TNIK have small molecule inhibitors targeting their kinase activities, and PIK3CA inhibitors have not produced a meaningful clinical response to date (*11–14*). In addition to the 3q chromosome, HNSCC is associated with amplification of 8q24 which includes *c-MYC* (*15, 16*). LZK phosphorylates and stabilizes the E3 ubiquitin ligase TRIM25, which ubiquitinates FBXW7, a subunit of the SKP1-Cullin-F-Box (SCF) complex that targets c-MYC for degradation (*17*). Loss of LZK-mediated TRIM25 phosphorylation, through depletion of LZK, leads to the degradation of the ligase, stabilization of FBXW7, and degradation of c-MYC (*17*).

We previously identified LZK (*MAP3K13*) as an amplified driver gene in 3q amplicon-positive HNSCC (*1*). We established that copy number gain at the 3q locus correlated with both an increase in *MAP3K13* mRNA and LZK protein abundance (*1*). Knocking down LZK with siRNA or shRNA significantly reduced cell viability and proliferation in 3q amplicon-positive HNSCC cells (CAL33, BICR56, Detroit562, and BICR6), but not control immortalized diploid cells (oral keratinocytes, OKF6, or bronchial epithelial cells, BEAS-2B) or HNSCC cells lacking amplified LZK (MSK921 and BICR22) (*1*). Depletion of LZK by doxycycline (dox)-inducible shRNA profoundly impaired colony formation in cells with *MAP3K13* gains or amplifications. Re-expression of an shRNA-resistant LZK in HNSCC cells fully or partially rescued the effect of LZK knockdown on cell density, confirming that these effects were specific to LZK depletion. Cells exhibiting impaired colony formation also exhibited a reduction in tumor burden in a xenograft mouse model of HNSCC, establishing an essential role for LZK expression in maintaining HNSCC cell viability (*1*).

There are currently no reported LZK inhibitors. However, there are potent ATP-competitive inhibitors of the related kinase DLK (*18*). Based on the greater than 90% homology in the kinase domains of LZK and DLK, here we tested the use of a DLK inhibitor as an inhibitor of LZK and this effective LZK inhibitor was then assessed for the ability to reduce HNSCC cell viability and suppress tumor growth *in vivo*. We defined a mechanism by which inhibition of LZK catalytic activity reduces HNSCC cell viability through destabilization of c-MYC. We found a kinase-independent mechanism by which LZK regulates GOF-p53, which led us to design PROTACs to target LZK for degradation. PROTACs are bifunctional molecules: One part binds to an E3 ubiquitin ligase and the second part or “warhead,” binds the target, in this case LZK. Effective PROTACs induce ubiquitination of the target by E3 ubiquitin ligase for subsequent proteasomal degradation. Keys to the success of a PROTAC are the affinity and specificity of the warhead for the target, effective recruitment of the E3 ubiquitin ligase and ternary complex formation, as well as suitable pharmacodynamic properties. Our lead LZK-targeting PROTAC abolished the kinase-dependent (c-MYC stabilization) and kinase-independent (GOF-p53 stabilization) oncogenic signaling mechanisms downstream of LZK. Our results demonstrate the promise of targeting LZK as a potential new treatment strategy for HNSCC patients.

## RESULTS

### Inhibition of LZK catalytic activity with GNE-3511 impairs viability and oncogenic phenotypes of HNSCC cells with amplified *MAP3K13*

To address if LZK is a favorable drug target for HNSCC, we used a potent DLK inhibitor, GNE-3511, that also inhibits the catalytic activity of LZK (*18*) (Figure 1A). We confirmed the high thermodynamic interaction affinity of GNE-3511 to LZK (Kd = 2.7 nM) using an ATP-independent competition binding assay (Figure 1B). LZK is a MAP3K [mitogen-activated protein (MAP) kinase kinase kinase] that directly phosphorylates the MAP2Ks (MAP kinase kinases) MKK7 and MKK4, leading to JNK pathway activation (*1, 19*). Exogenously overexpressed LZK activates the JNK pathway, which can be used as a readout to assess catalytic inhibition of LZK in cells (*1*). To verify that GNE-3511 inhibits LZK catalytic activity in cells, we induced expression of dox-inducible LZK in the 3q amplicon-positive HNSCC cell line CAL33 (Figure S1A). GNE-3511 potently inhibited LZK activity in cells (Figure 1C-F). Treatment of HNSCC cells harboring amplified *MAP3K13* (CAL33, BICR56, and Detroit562) with GNE-3511 resulted in an 80% or greater reduction in colony formation in the CAL33 and BICR56 cell lines and a 46% reduction in colony formation in Detroit562 cell line (Figure 1G, H), phenocopying the effect of LZK depletion (*1*). GNE-3511 produced only a minor or no reduction in colony formation in HNSCC cells lacking amplified *MAP3K13* (MSK921 and BICR22) (Figure 1G, H) and in BEAS2B cells, an immortalized and non-tumorigenic human lung epithelial cell line (Figure S1B).

**Figure 1.**
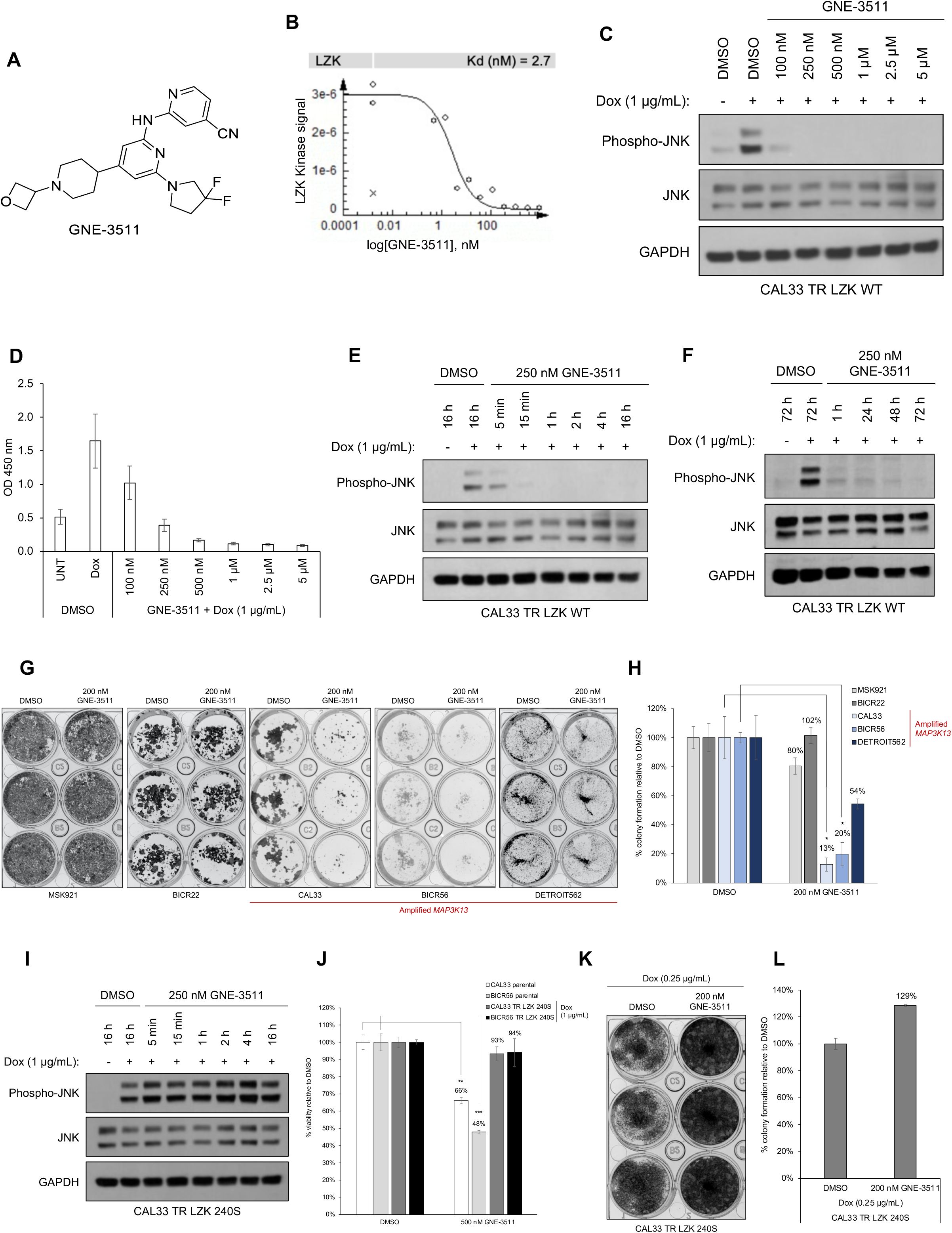
GNE-3511 induces LZK-dependent inhibition of cell viability and colony formation. **A.** Chemical structure of GNE-3511. **B.** Dose curve of the *in vitro* binding affinity of GNE-3511 to LZK assessed by Eurofin’s KdELECT KINOMEscan^TM^ profiling. The amount of kinase measured by qPCR (Signal; y-axis) is plotted against the corresponding compound concentration in nM in log10 scale (x-axis). Data points marked with an “x” were not used for Kd determination. **C.** Western blot showing the effect of increasing concentrations of GNE-3511 on JNK phosphorylation in response to doxycycline (dox)-induced expression of wild-type (WT) LZK in CAL33 cells. GAPDH served as the loading control. Data are representative of 3 independent experiments. **D.** Phospho-JNK ELISA analysis showing the effect of increasing concentrations of GNE-3511 on pJNK levels in response to dox-induced LZK expression in CAL33 cells. UNT, exposed to DMSO only. Data are shown as mean ± standard deviation for 3 experiments with triplicates each. **E, F.** Western blots showing time courses of the effect of GNE-3511 on JNK phosphorylation in response to dox-induced LZK WT expression in CAL33 cells. GAPDH served as the loading control. Data are representative of 3 independent experiments. **G, H.** Effect of GNE-3511 on colony formation by HNSCC cell lines with or without amplified *MAP3K13*. Panel G shows a representative experiment; panel H shows quantitative data for 3 experiments presented as the mean ± SEM. Significant differences were determined by Student’s *t-*test (*, p < 0.05). **I.** Western blot showing a time course of the effect of GNE-3511 on JNK phosphorylation in response to dox-induced expression of LZK with the drug-resistant Q240S mutation (TR LZK Q240S) in CAL33 cells. GAPDH served as the loading control. Data are representative of 3 independent experiments. **J.** Effect of GNE-3511 on cell viability in cells with or without overexpression of LZK Q240S. Expression of LZK Q240S was induced with dox. Viability was assessed 72 hours after addition of GNE-3511 or DMSO (as the vehicle control) and determined by MTS assay. Data are shown as mean ± SEM for 3 experiments with triplicates each. Significant differences were determined by Student’s *t*-test (**, p < 0.01; ***, p < 0.001). **K, L.** Effect of GNE-3511 on colony formation of CAL33 cells with dox-induced LZK Q240S. Panel K shows a representative experiment; panel L shows quantitative data for 3 experiments presented as the mean ± SEM.

Because kinase inhibitors are promiscuous compounds that often target multiple kinases and GNE-3511 was initially developed as a DLK inhibitor, we validated that the drug-induced decreases in cell viability were due to LZK inhibition. We generated and induced the expression of a drug-resistant mutant form of LZK (LZK^Q240S^) (Figure S1A, C), which maintains its catalytic activity in the presence of GNE-3511 (Figure 1I; Figure S1D). This mutant may have increased stability as its expression levels and activity are marginally increased compared to WT LZK (Figure S1C). When expressed in cells, LZK^Q240S^ prevented GNE-3511-induced decreases in viability of CAL33 and BICR56 cells (Figure 1J), and blocked GNE-3511-induced suppression of colony formation in CAL33 cells (Figure 1K, L). On the other hand, WT LZK expressing cells were more sensitive to GNE-3511, compared to CAL33 Q240S cells (Figure S1E). Thus, we confirmed that drug-induced decreases in cell viability are specifically due to inhibition of LZK.

### LZK enables oncogenic signaling through c-MYC in HNSCC

To define the mechanism by which catalytic activity of LZK maintains HNSCC cell survival, we first assessed the abundance of GOF-p53, a downstream target of LZK based on knockdown experiments (*1*). In contrast to LZK knockdown, GNE-3511 did not alter GOF-p53 abundance in CAL33 cells with the R175H GOF mutation in p53 or in BICR56, Detroit562, and MSK921 cells (Figure 2A, B), indicating that LZK regulates GOF-p53 in a kinase-independent manner. Instead, co-immunoprecipitation experiments indicated that LZK interacts with p53 (Figure 2C and S2A, B) and that the C-terminal acidic region of LZK is involved in this interaction (Figure S2A, B). We confirmed that LZK kinase activity was unnecessary for the interaction with p53 by coimmunoprecipitating p53 with the kinase-dead mutant LZK^K195M^ (Figure 2C; Figure S2B). Cell fractionation and immunofluorescence experiments suggested that LZK binds GOF-p53 in the cytoplasm of the cells (Figure S2C, S2D).

**Figure 2.**
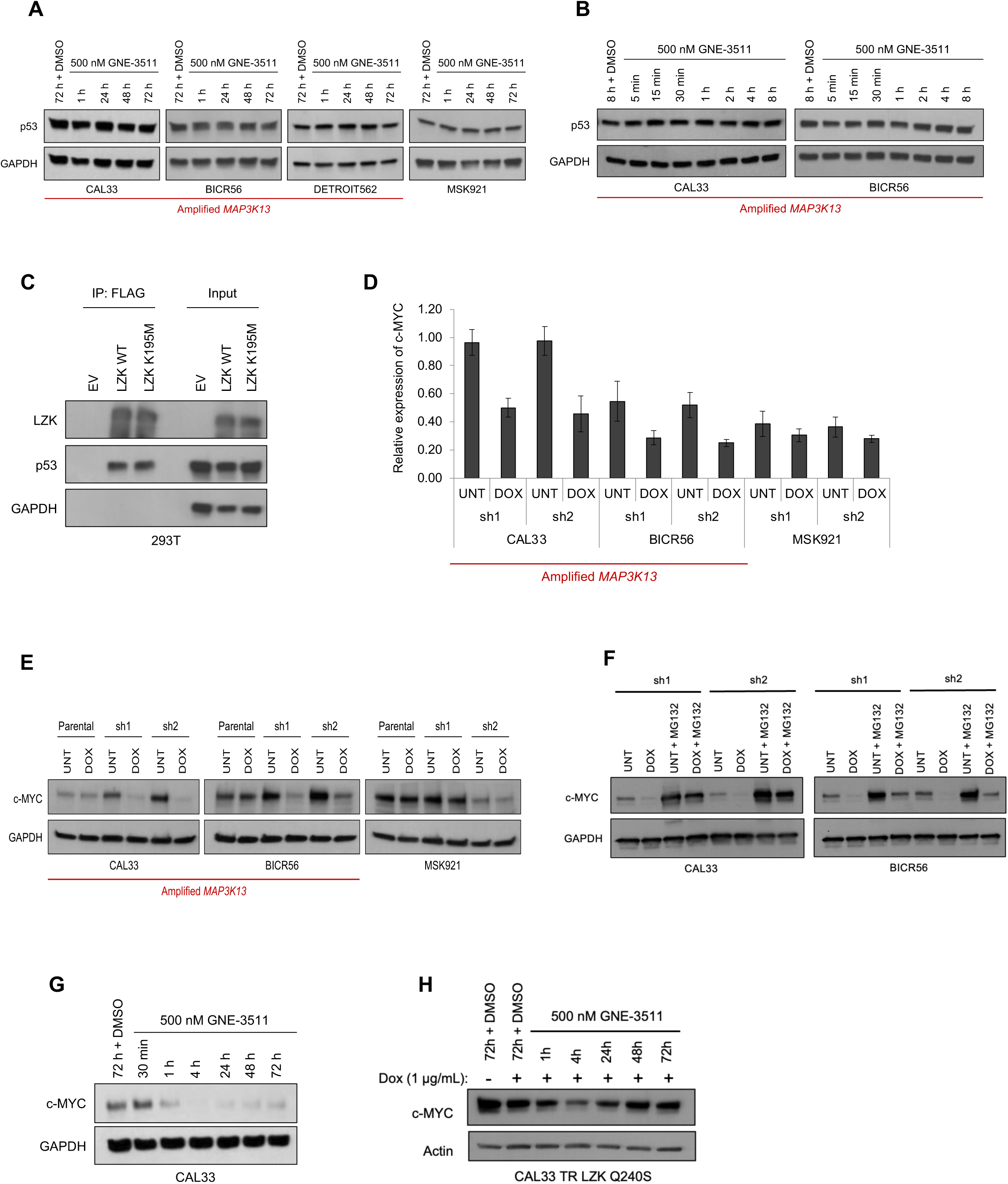
Kinase-dependent and - independent roles of LZK affect c-MYC and p53 abundance, respectively. A, **B.** Western blot showing p53 abundance in the indicated cells exposed to increasing times (5 minutes to up to 72h) of GNE-3511. GAPDH served as the loading control. Data are representative of 3 independent experiments. **C.** Coimmunoprecipitation of p53 with FLAG-tagged LZK from 293T cells detected by Western blot. GAPDH served as negative control for interaction and loading control. Data are representative of 3 independent experiments. EV, empty vector; LZK WT, wild-type FLAG-tagged LZK; LZK K195M, Flag-tagged kinase-dead K195M mutant LZK; IP, immunoprecipitation; Input, cell lysate. **D.** Normalized abundance of *c-MYC* in the indicated cell lines with dox-induced expression of silencing RNAs (sh1, sh2) targeting LZK. C-MYC abundance was determined by RPPA assay. Data for 3 experiments are presented as the mean ± SEM. **E.** Western blot showing the effect of dox-induced LZK knockdown (sh1, sh2) on c-MYC abundance in the indicated cell lines. Parental indicates the parental cell line without inducible shRNA. GAPDH served as the loading control. Data are representative of 3 independent experiments. UNT, no doxycycline. **F.** Western blot showing the effect of proteasome inhibition on c-MYC abundance in response to knockdown of LZK. The indicated cells with inducible LZK-targeted shRNA were untreated (UNT), exposed to doxycycline (DOX), or exposed to doxycycline and the proteasome inhibitor MG132 (10 µM). GAPDH served as the loading control. Data are representative of 3 independent experiments. **G, H.** Western blots showing time courses of the effect of GNE-3511 on CAL33 cells (G) or CAL33 cells with dox-induced expression of LZK Q240S (H). GAPDH served as the loading control. Data are representative of 3 independent experiments.

To identify other potential pathways LZK regulates, we performed a reverse phase protein array (RPPA), which identified c-MYC. Dox-inducible depletion of LZK with two unique LZK shRNAs (as described in Edwards *et al.* and Figure S2E) (*1*) reduced c-MYC abundance in 3q amplicon-positive HNSCC cells (CAL33 and BICR56) but not control cells (MSK921) (Figure 2D, E). The decrease in c-MYC induced by knockdown of LZK depended on proteasome-mediated degradation; addition of the proteasome inhibitor MG132 suppressed this reduction (Figure 2F). To determine if LZK catalytic inhibition reduced c-MYC abundance, we treated CAL33 cells with GNE-3511 at 500 nM, which reduced c-MYC abundance, that persisted for 72 hours (Figure 2G). The drug-resistant mutant LZK^Q240S^ blocked the persistent inhibitor-induced reduction in c-MYC abundance, indicating that LZK catalytic activity is essential for maintaining c-MYC abundance (Figure 2H). Thus, our data showed that LZK has both kinase-dependent activity (related to c-MYC) and kinase-independent functions (related to GOF-p53) that promote cancer.

### Blocking LZK function delays cell cycle progression and induces quiescence

c-MYC activity promotes progression through the cell cycle (*20*), therefore we assessed the impact of GNE-3511 on the cell cycle in the 3q amplicon-positive cell lines CAL33 (Figure 3) and SCC15 (Figure S3) (*21, 22*). GNE-3511 treatment caused a shift in DNA content distribution (Figure 3A), and a decrease in EdU incorporation in CAL33 cells (Figure 3B). Single-cell scatter plots of both DNA content and EdU incorporation also showed a change in the cell cycle distribution in cells treated with as low as 100 nM GNE-3511 (Figure 3C-D). Single-cell assay tracking asynchronously dividing cells further showed that GNE-3511 delayed the increase in CDK2 activity (Figure 3E-3H; Figure S3A-F) and exhibited dose-dependent effects in delaying mitosis (Figure 3I). The sustained reduction in CDK2 activity in response to GNE-3511 (Figure 3G-H; Figure S2F) suggested that the cells experienced a G2 arrest (*22*). Furthermore, quantification of the single-cell data showed a significant increase in the percentage of cells halting in G2 phase (Figure S3G, S3H). To determine if apoptosis also contributed to the decline in cells undergoing mitosis, we examined the cultures for apoptotic cells. Even after 54 hours of exposure to the inhibitor, we rarely observed apoptotic cells (Figure 3J), and no cleavage of PARP or caspase-3 pro-apoptotic markers were detected in cells treated with the inhibitor for up to 96h (Figure S4A, B), suggesting that the delay in progression through the cell cycle resulted primarily from an increase in cells undergoing cell cycle arrest.

**Figure 3.**
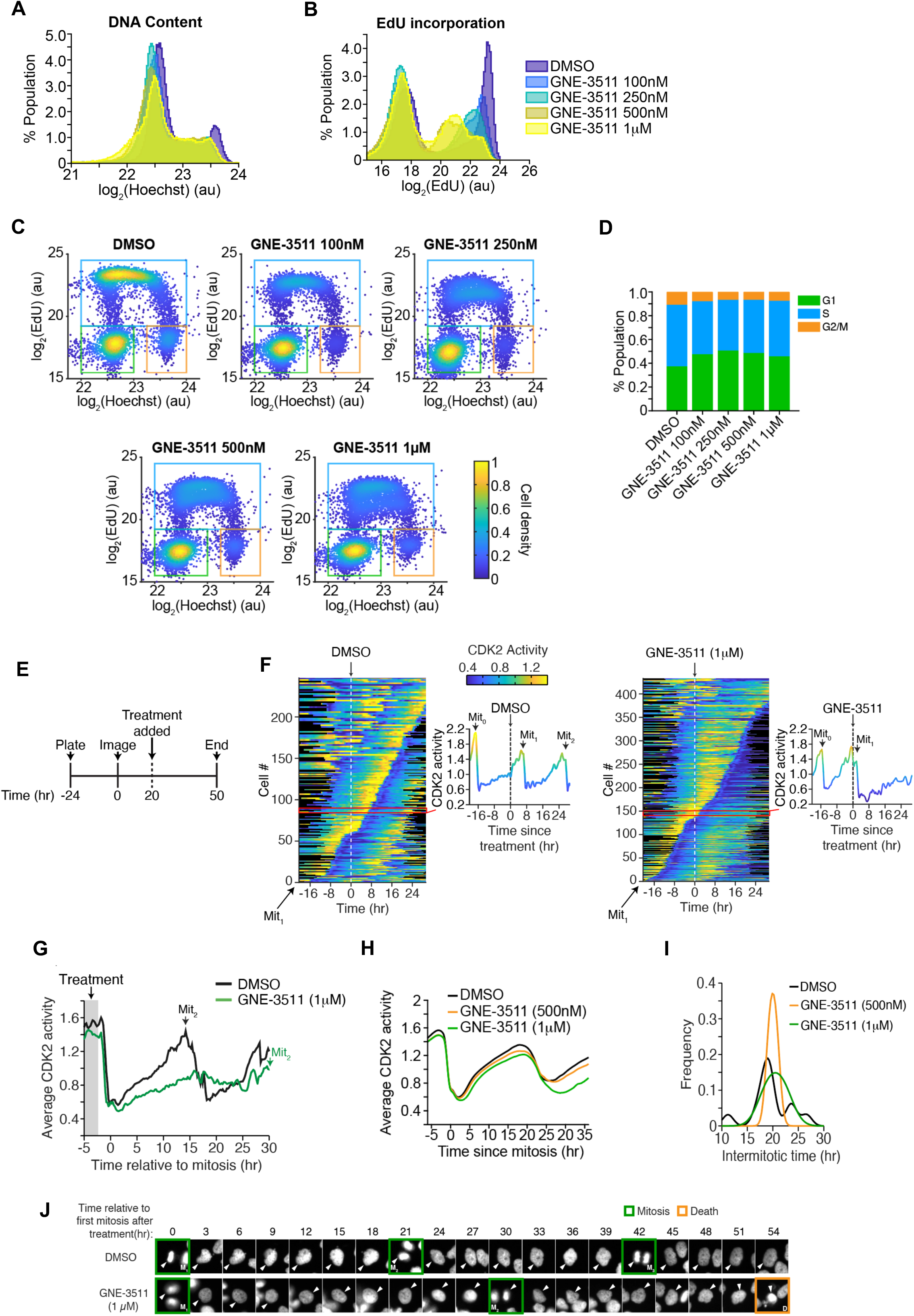
Inhibition of LZK with GNE-3511 impairs progression through the cell cycle. **A.** Histograms of DNA content for CAL33 cells treated with the indicated concentration of GNE-3511 for 48hr. Data are representative of 3 independent experiments. **B**. Histograms of EdU incorporation for CAL33 cells treated with the indicated concentration of GNE-3511 for 48 hr. Cells were treated for 48 hr and then grown in media containing EdU for 30 min prior to fixation. Data are representative of 3 independent experiments. **C.** Density scatter plot of the DNA content and EdU incorporation in single-cells after treatment with the indicated concentration of GNE-3511 for 48 hr. Colored boxes represent different cell cycle phases (Green, G1; Cyan, S; Orange, G2/M). Data are representative of 3 independent experiments. **D.** Bar graph of the distribution of cell cycle phases from (C). **E.** Diagram of experimental protocol in CAL33 cells. Mitotic events were monitored using the peaks of CDK2 activity in each cell. Images were acquired 20 hours before exposure to treatment (DMSO or GNE-3511) and then every 12 minutes for an additional 30 hours. **F.** Cell cycle progression monitored by the activity of CDK2 in individual asynchronously growing CAL33 cells exposed to DMSO (vehicle control) or GNE-3511. Individual cell example traces, indicated by a red line, are shown to the right of each heatmap. Mitotic events are noted as Mit_0_ (before exposure), Mit_1_ (first division after exposure), Mit_2_ (second division after exposure). Data are representative of 3 independent experiments. **G.** Effect of GNE-3511 on the average time between the first mitotic event after exposure and the second mitotic event (Mit_2_) in CAL33 cells. Data are representative of 3 independent experiments. **H.** Graph showing the effect of DMSO or GNE-3511 on average CDK2 activity in CAL33 cells during progression through the cell cycle. **I.** Frequency histogram showing the concentration-dependent effect of GNE-3511 on intermitotic time of CAL33 cells. Data are representative of 3 independent experiments. **J.** Microscopy film strips show three individual cells every three hours over a total of 54 hours after treatment with DMSO or GNE-3511, with green boxes outlining mitotic cells and orange boxes outlining dying cells.

### Targeting LZK is effective for 3q-amplified HNSCC tumors in vivo

We identified 5 patient-derived xenograft mouse models (PDX 391396-364-R, 295223-140-R, 899872-116-R, 226611-082-R, and 653398-132-R) with amplified *MAP3K13* (at least 4 copies) and 31 PDXs with gains of *MAPK3K13* (3 copies) in 58 HNSCC PDX models from the NCI Patient-Derived Models Repository (PDMR) (Figure 4A, B) (*23–25*). *MAP3K13* was a top gene amplified within chromosome 3 in these patient samples (Figure 4A, B). Furthermore, increased copy number of *MAP3K13* was associated with an increase in mRNA abundance (Figure 4C). We selected 2 PDX models with *MAP3K13* amplification and high mRNA expression (391396-364-R and 295223-140-R) and 2 PDX models without amplification and with low *MAP3K13* mRNA expression (328373-195-R and 959717-210-R) (Supplementary table S1) and tested the effect of GNE-3511 *in vivo*. GNE-3511 reduced tumor growth for both *MAP3K13*-amplified PDX models (Figure 4D) and had little effect on the 2 PDX models without increased *MAP3K13* expression (Figure 4E). None of the mice exhibited toxicity based on stable body weight throughout the treatment period (Figure S5A, B). For PDX 391396-364-R, 3 mice had tumor regression such that tumors were undetectable. Overall, these results indicated that inhibiting the catalytic activity of LZK has therapeutic potential for 3q-amplicon positive HNSCC patients.

**Figure 4.**
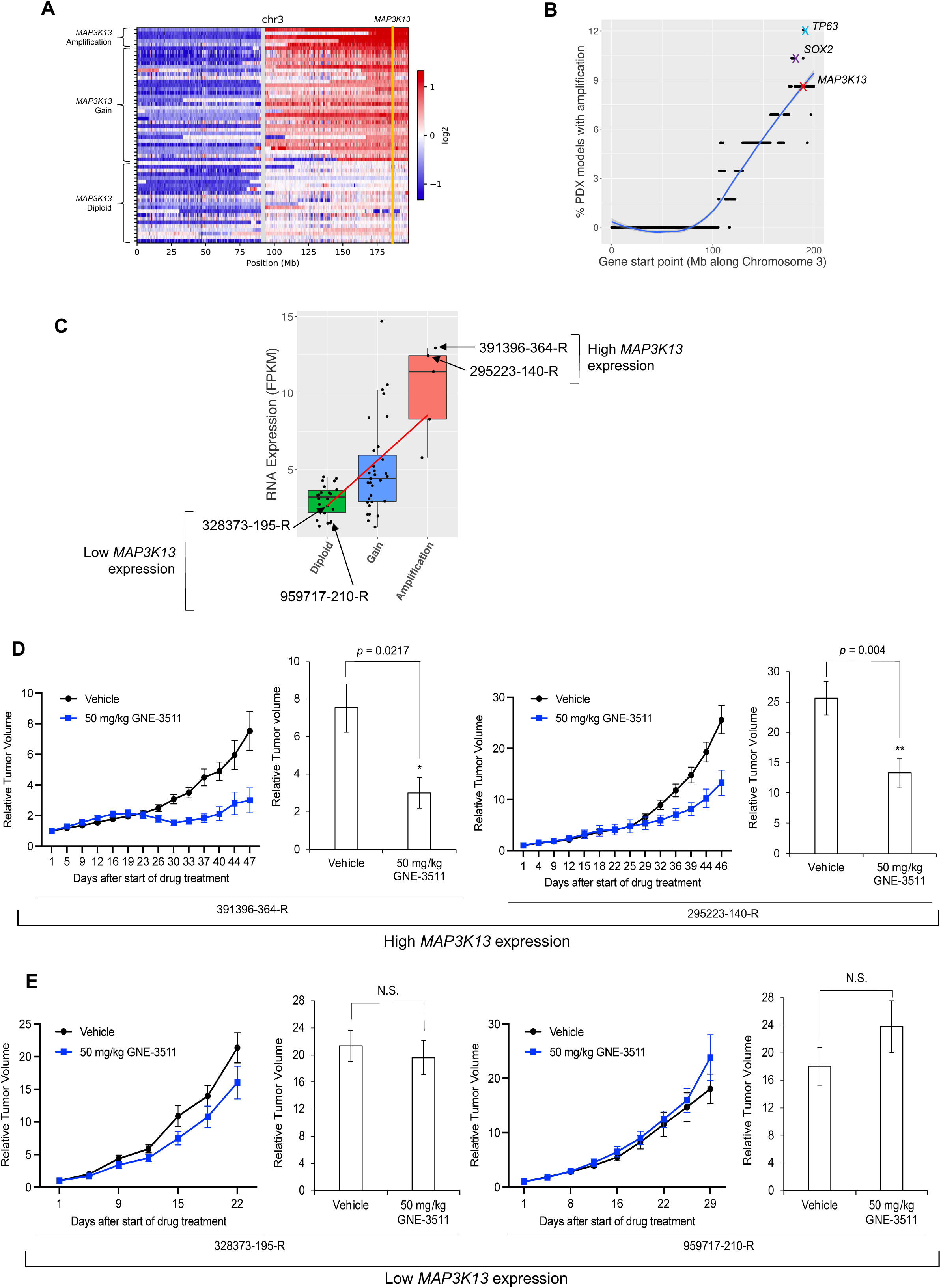
GNE-3511 impairs growth of *MAP3K13*-amplified PDX tumors in mice. **A.** Copy number (CN) profiles of fifty-eight HNSCC PDX mouse models on chromosome 3 obtained from the NCI PDMR. Each row indicates the copy number profile of one PDX model. Models were ordered by *MAP3K13* copy number data (highlighted as yellow line). Heatmap color indicates the log2 ratio of copy numbers. **B.** Graph represents the percentage of the HNSCC PDX models with amplification of each gene on chromosome 3. Genes were ordered by gene start point along chromosome 3. *MAP3K13* is marked with a red cross. Blue line is the regression line by loss method. The 3 genes with the highest frequency of amplification are labelled. **C.** Abundance of transcripts for *MAP3K13* in average fragments per kilobase million (FPKM) in PDX models diploid for MAP3K13 (= 2 copies), with gain of *MAP3K13* (>2 and < 5 copies), or amplification of *MAP3K13* (≥ 5 copies) (based on the clustering in C). Each PDX is shown as a dot and box. Copy number of *MAP3K13* is highly correlated with gene expression (ANOVA, *p* = 1.34e^-6^). The 4 PDX models used for subsequence study are indicated. **D, E.** Effect of GNE-3511 on tumor volume in 2 PDX models with *MAP3K13* amplification and high expression (F, 10 mice per treatment condition) and on 2 PDX models with diploid *MAP3K13* and low expression (G, 10 mice per treatment condition). Mice were administered the indicated treatments (50 mg/kg, q.d., five days on/two days off). Mean relative tumor volumes ± SEM are shown. Average relative tumor volume at the end of treatment Mean ± SEM is shown to the right of the graphs. Statistically significant differences were assessed by Student’s *t*-test; (\*\**p* < 0.01, \**p* < 0.05; N.S., not significant).

### An LZK-targeted PROTAC reduces both the kinase-dependent and - independent cancer-enabling activities of LZK

One way to block all pro-cancer LZK-mediated pathways is to eliminate LZK protein using LZK-targeted PROTACs (*26–28*). Such an approach should reduce both c-MYC and GOF-p53 (Figure 5A). GNE-3511 is unsuitable for PROTAC development due to the oxetane moiety that prevents attachment of the linker. Therefore, to develop LZK-targeting PROTACs, we tethered E3-ligase-binding warheads to another DLK inhibitor compound #21 (Figure S6A) (*18*) using various linkers. Compound #21 has a structural similarity to GNE-3511, in which the precursor amine of GNE-3511 is functionalized with an amide rather than an oxetane, making it a more favorable moiety for the addition of the linker and E3 ligase ligand. Compound #21 has a comparable binding affinity towards LZK as GNE-3511 (Kd = 8.9 nM), as assessed using an ATP-independent competition binding assay (Figure S6B). Compound #21 impaired LZK-induced JNK phosphorylation with a potency and duration of inhibition similar to that of GNE-3511 (Figure S6C-S6F). Like GNE-3511, compound #21 impaired colony formation of 3q amplicon-positive HNSCC cells (Figure S6G, S6H). Furthermore, the inhibitory effects were substantially reduced or absent in cells expressing LZK^Q240S^ (Figure S6I, S6J), confirming that the effects depended on LZK.

**Figure 5.**
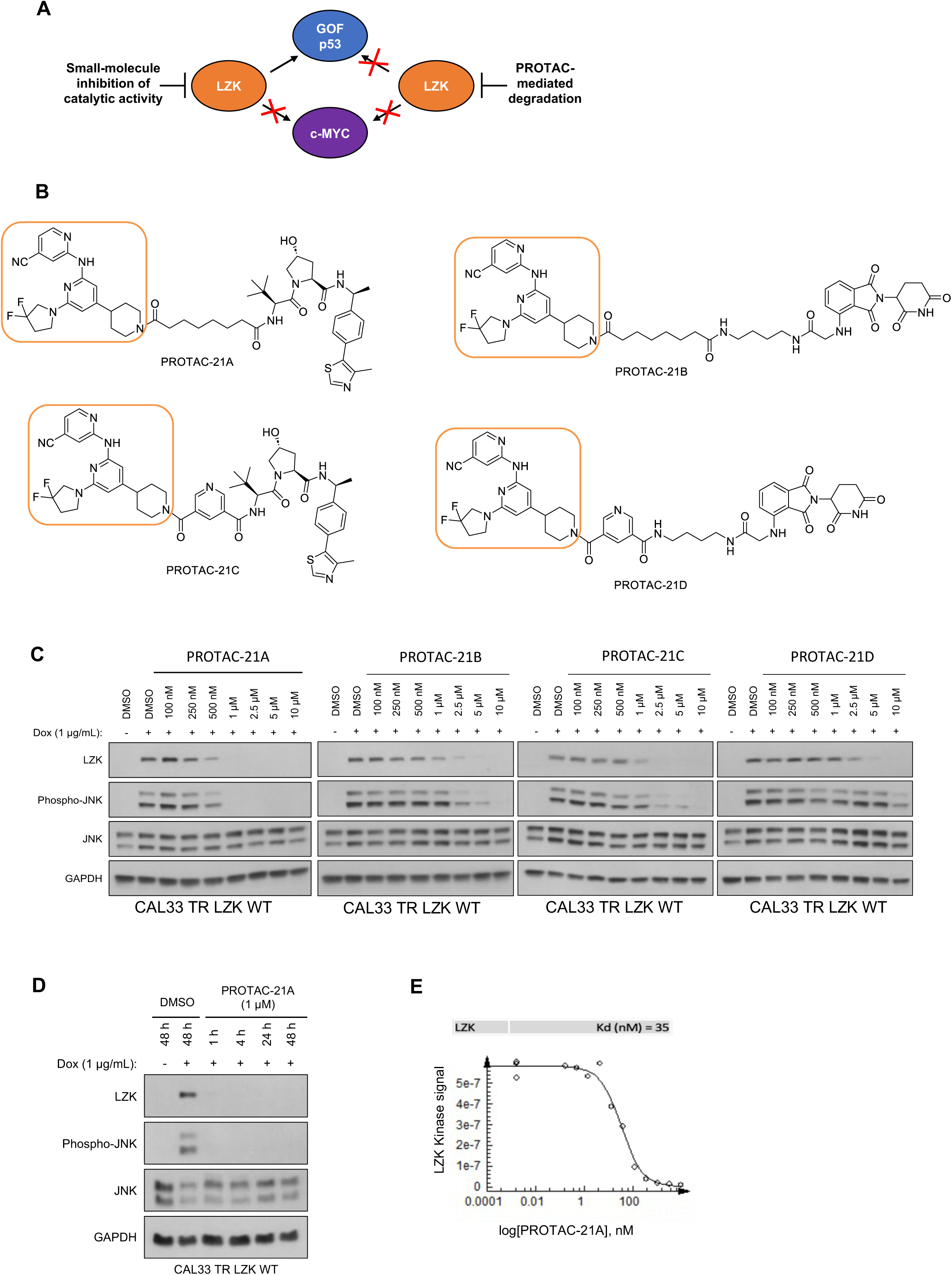
Development of LZK-targeting PROTACs with compound 21 as the pharmacophore. **A.** Diagram showing how PROTAC-mediated degradation acts to target both kinase-dependent (c-MYC) and - independent (GOF-p53) pro-tumor functions of LZK. **B.** Chemical structures of PROTAC-21A, PROTAC-21B, PROTAC-21C, and PROTAC-21D. The compound #21 warhead is outlined in orange. **C.** Western blots showing the effect of increasing concentrations of PROTAC-21A, PROTAC-21B, PROTAC-21C, and PROTAC-21D on LZK expression and JNK phosphorylation in response to dox-induced expression of LZK in CAL33 cells. GAPDH served as the loading control. Data are representative of 3 independent experiments. **D.** Western blots showing a time course of the effect of PROTAC-21A on LZK expression and JNK phosphorylation in response to dox-induced LZK WT expression in CAL33 cells. GAPDH served as the loading control. Data are representative of 3 independent experiments. **E.** Dose curve of the *in vitro* binding affinity of PROTAC-21A to LZK assessed by Eurofin’s KdELECT KINOMEscan^TM^ profiling. The amount of kinase measured by qPCR (Signal; y-axis) is plotted against the corresponding compound concentration in nM in log10 scale (x-axis).

Our lead PROTAC-21A (Figure 5B) had similar effects to those of compound #21: dose-dependent, persistent inhibition of LZK-induced JNK phosphorylation that was maintained out to 48 hrs (Figure 5C, D), with an LZK binding affinity of 35 nM (Figure 5E). Additional PROTACs generated using compound #21, PROTAC-21B, PROTAC-21C and PROTAC-21D (Figure 5B) also promoted degradation of LZK at higher concentrations (Figure 5C). We confirmed that PROTAC-21A reduced LZK abundance through a mechanism that involved the ubiquitin-like molecule NEDD8 (Figure 6A) and the proteasome (Figure 6A, B). Interestingly, in cells treated with MG132 there is still some residual JNK phosphorylation, compared to cells treated with MLN4924 (Figure 6A). To assess if the increase in JNK phosphorylation is due to MG132 treatment, we treated cells with MG132 alone or in combination with PROTAC 21A and observed an increase in JNK phosphorylation in cells treated with MG132, indicating this increase is likely due to a stress response leading to JNK activation (Figure S7). Evaluation of kinome selectivity against a broad panel of >450 kinases revealed that both compound#21 and PROTAC-21A are selective for DLK/LZK (Figure S8 and Data File S1). We further validated the selectivity of PROTAC-21A using a mass spectrometry-based global proteomic analysis in CAL33 cells upon induction of LZK expression and treatment with DMSO control or 1 µM PROTAC-21A (Figure S9A and S9B) and showed that LZK along with 10 other proteins were downregulated upon PROTAC treatment (Figure S9B, Data File S2). Notably, 10 out of the 11 downregulated proteins by PROTAC-21A were upregulated by dox-induced expression of LZK, indicative of being downstream effectors of LZK (Figure S9C, Data File S2). PROTAC-21A also reduced colony formation of 3q amplicon-positive HNSCC cells (Figure 6C, D). However, much higher concentrations were required than for the small molecule inhibitors. Reduction of phosphorylated JNK (detected by Western blot) required 1 µM PROTAC-21A (Figure 5C, D), whereas GNE-3511 or compound #21 achieved a similar level of reduction at 250 nM (Figure 1C, S6E). At 1 µM, PROTAC-21A reduced colony forming ability by amplicon-positive CAL33 and BICR56 cells without having any major effect on BICR22 and BEAS-2B control cell lines (Figure 6C, D). At higher concentration of 2.5 µM, PROTAC-21A completely abrogated colony formation in CAL33 and BICR56 cells; however, some off-target toxicity was observed in control BICR22 and BEAS-2B cells (Figure S10A). We then showed that the cis-epimer of PROTAC-21A (Figure S10B) that inhibits LZK activity but does not degrade it (Figure 6E) does not have any significant effect on colony forming ability by the HNSCC cell panel at 1 µM (Figure 6F). Permeability analysis showed that PROTAC-21A and PROTAC-21C had limited cell permeability (Figure S10C), and pharmacokinetic *in vivo* studies of the PROTACs demonstrated variable tissue distribution (Figure S10D), providing a rationale for the higher concentration and lower effectiveness of PROTAC-21A than the small molecule LZK inhibitors and the need for optimization to develop better drug-like compounds.

**Figure 6.**
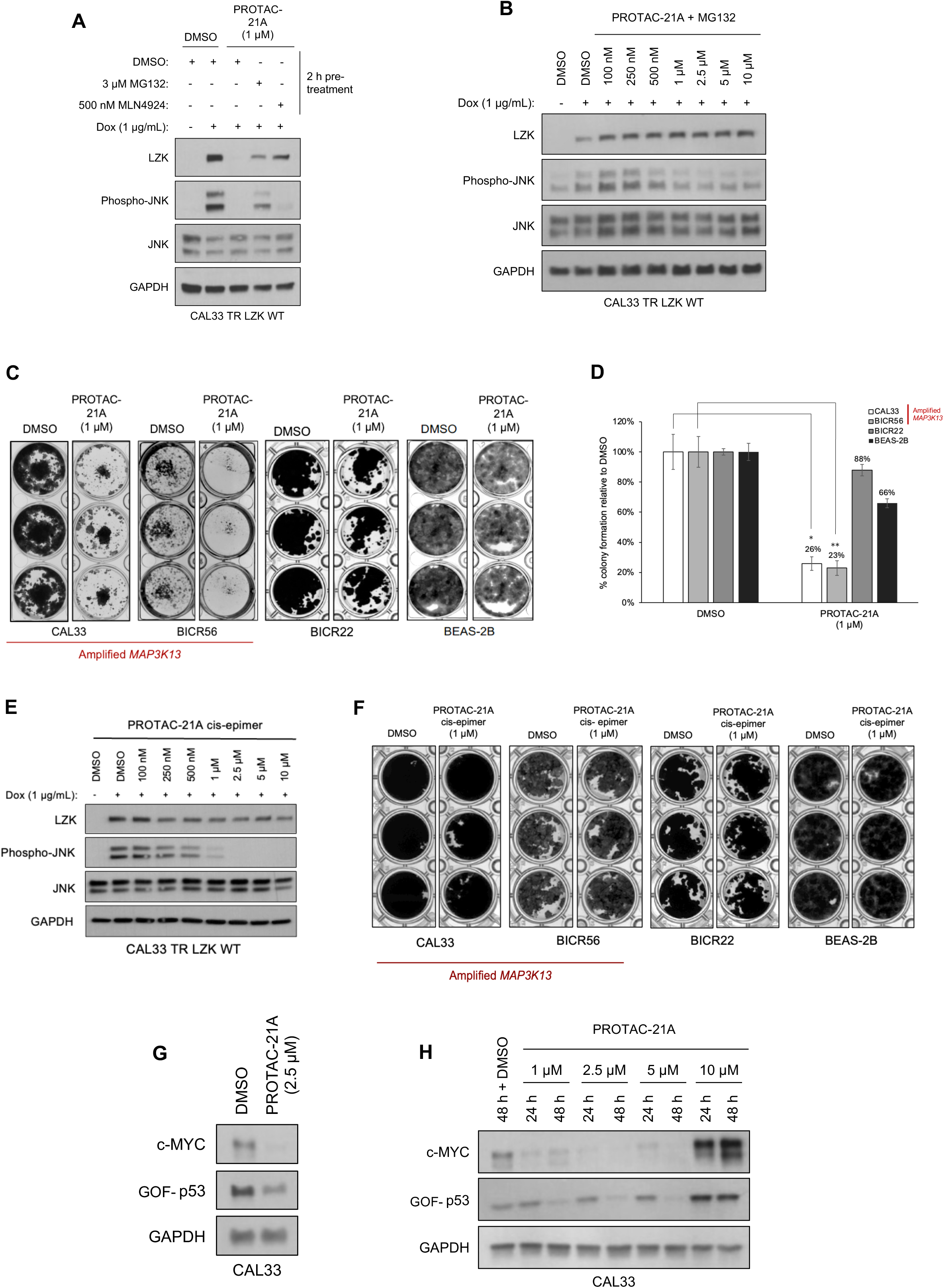
Lead PROTAC-21A induces proteasome-mediated degradation of LZK, targets c-MYC and GOF-p53, and decreases HNSCC viability. **A.** Western blot showing dependence of PROTAC-21A-mediated degradation of LZK involves both ubiquitin-like molecule NEDD8 and the proteasome. LZK WT was induced with dox in CAL33 cells and the cells were pre-treated for 2 hours with either MG132 to inhibit the proteasome or MLN4924 to inhibit NEDD8 and then exposed to the indicated concentration of PROTAC-21A. Expression levels of LZK and phosphorylated JNK were monitored. GAPDH served as the loading control. Data are representative of 3 independent experiments. **B.** Western blot showing LZK and phosphorylated JNK levels in dox-induced CAL33 LZK WT cells exposed to 10 µM MG132 and increasing concentrations of PROTAC-21A. Data are representative of 3 independent experiments. **C, D.** Effect of PROTAC-21A on colony formation by the indicated cell lines. DMSO served as the vehicle control. Panel C shows a representative experiment; panel D shows quantitative data for 3 experiments presented as the mean ± SEM. Significant differences were determined by Student *t-*test (**, p < 0.01; ***, p < 0.001). **E.** Western blots showing the effect of increasing concentrations of PROTAC-21A enantiomer on LZK expression and JNK phosphorylation in response to dox-induced expression of LZK in CAL33 cells. GAPDH served as the loading control. Data are representative of 3 independent experiments. **F.** Effect of PROTAC-21A enantiomer on colony formation by the indicated cell lines. DMSO served as the vehicle control. Data shows a representative experiment. **G, H.** Western blot showing the effect of PROTAC-21A on gain-of-function p53 and c-MYC abundance in CAL33 cells. GAPDH served as the loading control. Data are representative of 3 independent experiments.

To establish that LZK degradation had the predicted effects on GOF-p53 and c-MYC, we showed that addition of PROTAC-21A to CAL33 cells reduced c-MYC abundance at both 24 hours and 48 hours and reduced GOF-p53 abundance at 48 hours (Figure 6G, H). However, the highest concentration of PROTAC-21A tested (10 µM) produced a paradoxical increase in both c-MYC and GOF-p53 compared to their abundance in the control. This may be due to the high concentration of the PROTAC having off-target effects that will promote the degradation of numerous kinases and other possible targets and block the function of LZK depletion to cause a decrease in c-MYC and GOF-p53 abundance. We aim to explore this interesting observation in future studies. Collectively, these data suggested that a PROTAC-based strategy is an option for abrogating both the kinase-dependent and - independent cancer-enabling activities of LZK but that additional optimization, especially regarding cell permeability, is required for application *in vivo*.

## DISCUSSION

*MAP3K13* is a driver gene that results in increased *MAP3K13* mRNA and LZK protein in 3q amplicon-positive HNSCC (*1*). GOF mutations in p53 are a common early occurrence in HNSCC and convert p53 from a tumor suppressor into a tumor promoter (*2, 29–32*). Previously, we determined that LZK is important for stabilizing GOF-p53 in HNSCC (*1*). Our previous work established that knocking down LZK reduced cell viability, proliferation, and colony formation of 3q amplicon-positive HNSCC cells and reduced tumor burden in a xenograft mouse model of HNSCC (*1*). Here, we determined that c-MYC is a second cancer-promoting target of LZK in HNSCC and that LZK is required to maintain c-MYC abundance. We also showed that an inhibitor of the related kinase DLK, GNE-3511, effectively and specifically impaired viability and colony formation of 3q amplicon-positive HNSCC cells and significantly decreased expression of c-MYC, which could be rescued by expression of a drug-resistant mutant form of LZK. The translational relevance of targeting LZK was shown through the inhibition of growth and even regression in HNSCC PDX models harboring amplified *MAPK3K13* compared to control models that are diploid for MAP3K13 (Figure 7).

**Figure 7.**
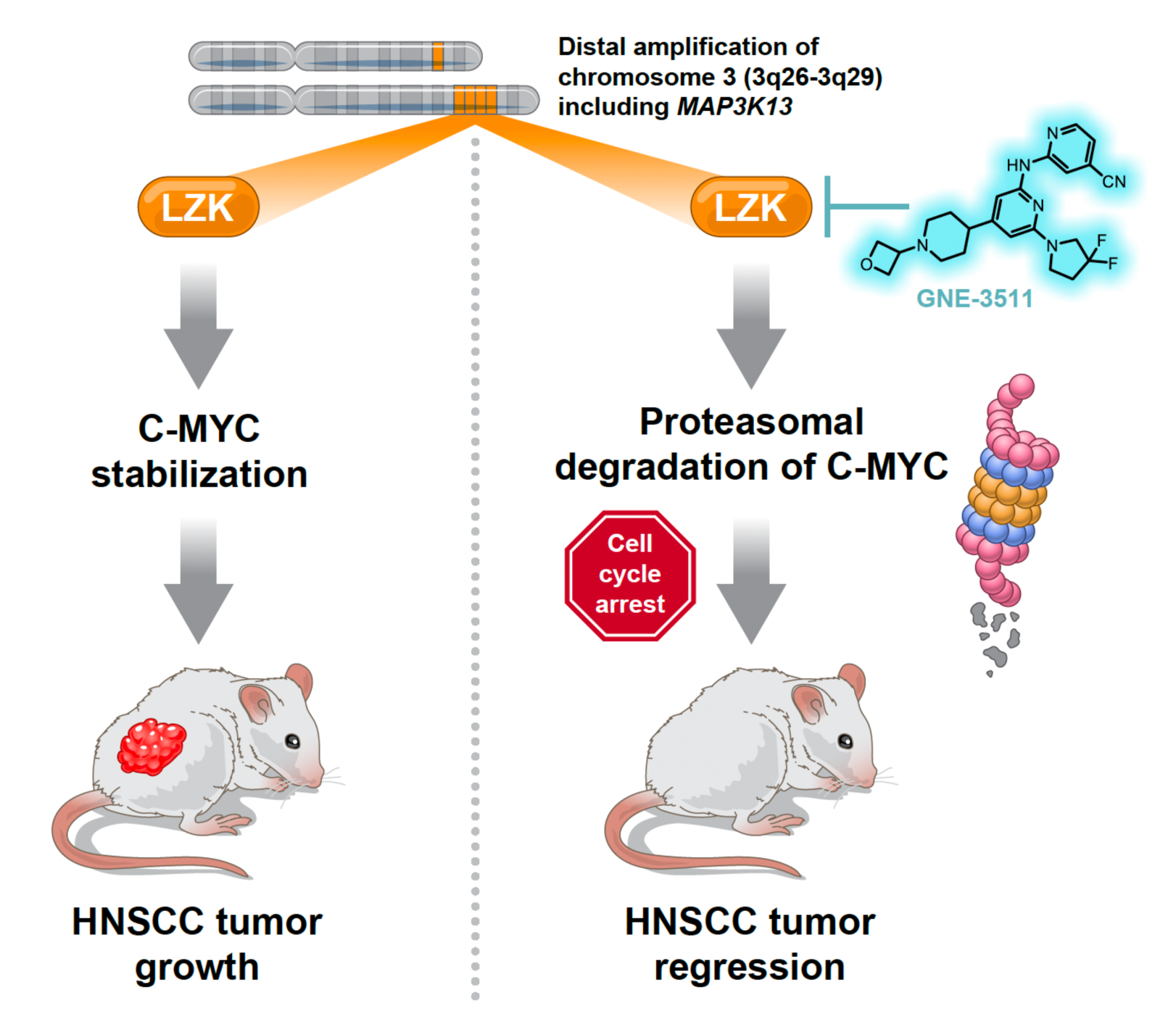
LZK inhibition suppresses Head and Neck Squamous Cell Carcinoma tumor growth via c-MYC. Amplified LZK regulates c-MYC stability in head and neck squamous cell carcinoma promoting tumor growth. Inhibition of LZK activity via GNE-3511 promotes proteasomal degradation of c-MYC causing cell cycle arrest and suppression of tumor growth in HNSCC.

In addition, we found that LZK-mediated stabilization of GOF-p53 was not inhibited by GNE-3511; however, stabilization of c-MYC was blocked by GNE-3511 in an LZK-dependent manner. Thus, maximal effectiveness in targeting LZK may require blocking both the kinase-dependent and kinase-independent actions of LZK. Optimizing LZK-targeting PROTACs will be extremely important going forward to achieve efficacy at low nanomolar concentrations as LZK depletion completely abrogates colony formation in HNSCC cell with amplified *MAP3K13*. In comparison, we observe a significant decrease in viability in cells treated with the LZK inhibitor, GNE-3511, however colonies still remain, highlighting the importance of the non-catalytic functions of LZK to promote cancer phenotypes.

Patients with recurrent or metastatic head and neck squamous cell carcinoma have limited effective treatment options. Pembrolizumab has recently been approved as a first-line therapy for this group (*33, 34*), yet response rates are still low, indicating a significant need for more effective treatments. Our research identifies a unique opportunity to develop a first-in-class LZK inhibitor for the approximately 70% of HNSCC patients that exhibit gains or amplifications in LZK. The promising *in vivo* results we have observed suggest that an optimized, orally bioavailable LZK inhibitor or PROTAC could serve as a viable therapeutic strategy. Additionally, the minimal expression of LZK in the human body implies that an LZK inhibitor would function as a precision medicine, specifically targeting tumor cells while minimizing toxic side effects. Squamous cell carcinomas (including HNSCC, ESCC, and LSCC) are unique cancers, primarily driven by copy number gains and losses, rather than drivers with gain-of-function mutations such as KRAS and EGFR (*2*). Recently, Grish and colleagues have compellingly demonstrated that amplified genes are potent oncogenes and represent a relatively untapped source of new therapeutic targets (*35*). These amplified drivers likely work together, and identifying co-amplified partners to target alongside LZK, such as PI3K, opens up an exciting area for future exploration, especially as combination therapies show improved efficacy in clinical settings (*36*). In summary, our study represents an important step in demonstrating that LZK-targeting inhibitors and degraders hold significant promise for treating a substantial subset of HNSCC patients. Future research will focus on developing potent LZK inhibitors and degraders that can be used alone or in combination as exciting new therapies for these patients.

## Methods

### Mice

NCI-Frederick is accredited by AAALAC International and follows the Public Health Service Policy for the Care and Use of Laboratory Animals. Animal care was provided in accordance with the procedures outlined in the “Guide for Care and Use of Laboratory Animals (National Research Council; 1996; National Academy Press; Washington, D.C.).

For our in-house patient-derived xenograft mouse experiment, we used six-to eight-week-old immunodeficient NOD-scid IL2R gamma^null^ (NSG) female mice obtained from the NCI Center for Cancer Research Animal Resource Program. All mice were maintained under pathogen-free conditions in the NCI-Frederick immunocompromised suite. Animal handling and experimental procedures were approved by the National Cancer Institute (NCI) Animal Care and Use Committee (ACUC) and was performed within the limits of a license granted by the Home Office according to the Animals (Scientific Procedures) Act 1986. For the drug studies, mice were randomly assigned experimental groups to prevent bias towards tumor size. After, an identification number was assigned to each animal and the researchers were not blinded to subsequent treatment and data collection.

Tumor pieces at approximately 2 × 2 × 2 mm^3^ from an HNSCC patient containing amplified or diploid *MAP3K13* were implanted subcutaneously with Matrigel (Corning) in the mice according to the SOP50101 Implantation and Cryopreservation of Tissue for PDX Generation protocol from the NIH Patient-Derived Models Repository (PDMR). Five NSG mice were used for initial implantation of the cryopreserved tumor fragments. Body weights and tumor size were measured twice weekly. The tumors were harvested when they reached approximately 1,000 mm^3^ and were used to generate the PDX mouse model to test GNE-3511. For the efficacy study, passage one of the fresh PDX tumor fragments were implanted into NSG mice using the protocol stated previously. Twenty NSG mice were used (10 for vehicle control and 10 for GNE-3511 treatment). Body weights and tumor sizes were measured twice weekly until tumors reached approximately 150–200 mm^3^, at which point the mice were randomly assigned to treatment cohorts with control or GNE-3511 for approximately 4–8 weeks. The study endpoints were over 20% body weight loss, tumor volume exceeding 2.0 cm^3^ in diameter, or significant (greater than 80%) tumor regression observed with treatment. The GNE-3511 was dissolved with 60% PEG 300 MW, 3 eq of 0.1 M HCl, saline (vehicle) and administered daily via intratumoral injection at 50 mg/kg. Body weights and tumor sizes were measured twice weekly after the start of drug administration. At the endpoint of each study, tumors were harvested, cleaned, weighed, and photographed for analysis.

For PK analysis of PROTAC-21A and PROTAC-21C, female NSG mice between the weight of 17-25 grams were obtained from The Jackson Laboratory. The PK animal study protocol was approved by ACUC of Division of Veterinary Resources at the National Institutes of Health (NIH).

The *in vitro* ADME properties of PROTAC-21A and PROTAC-21C were studied with the standard Tier I assays, i.e. kinetic solubility assay, parallel artificial membrane permeability assay (PAMPA) and rat liver microsomal (RLM) stability assay (*37–39*). The pharmacokinetics of PROTAC-21A and PROTAC-21C were determined in female NSG mice after a single intraperitoneal injection of 50 mg/kg. Mice used in this study (n = 3 per time point) were not fasted. PROTAC-21A and PROTAC-21C were dissolved with a formulation of 40% PEG 300 MW, 3 eq of 0.1 M HCl, saline. The dosing solutions were prepared fresh prior to the drug administration. The injection volume was 10 mL/kg. Blood samples were collected from each mouse before dosing (t = 0) and at 0.083, 0.25, 0.5, 1, 2, 4, 7 and 24 hour for PROTAC-21A; and at 0.083, 0.25, 0.5, 1, 2, 4, 7, 24 and 48 hour for PROTAC-21C. K_2_EDTA was used as the anti-coagulant. Blood samples was centrifuged at 14,000 rpm at 4 °C to obtain plasma. After blood collection, liver, kidney, lung, and brain tissues were collected, weighed and snap frozen with dry ice. All samples were stored at −80 °C until the analysis.

Drug concentrations in plasma and tissue samples were measured by a qualified UPLC-MS/MS method with a Waters Acquity I-Class UPLC interfaced with a Waters TQ-S mass spectrometer (Waters). The lower limit of quantitation (LLOQ) was 1 ng/mL for plasma and 1 ng/g tissue for liver, kidney, lung, and brain. PK parameters were calculated using the non-compartmental method of the pharmacokinetic software package Phoenix WinNonlin, version 6.2 (Certara). The area under the plasma concentration versus time curve (AUC) was calculated using the linear trapezoidal method. Where warranted, the slope of the apparent terminal phase was estimated by log-linear regression using at least 3 data points, and the terminal rate constant (λ) was derived from the slope. AUC_0-οο_ was estimated as the sum of the AUC_0-t_ (where t is the time of the last measurable concentration) and C_t_/λ. The apparent terminal half-life (t_½_) was calculated as 0.693/λ.

### Human Samples

Tumor fragments from HNSCC patients containing amplified or diploid *MAP3K13* were obtained from the NIH PDMR (Supplementary table S1).

### Bioinformatic analyses of HNSCC PDX mouse models of NCI PDMR

For nucleic acid extraction, library prep, whole-exome sequencing, and whole-transcriptome sequencing, please see the documents from the NIH PDMR SOPs. An in-house bioinformatics pipeline was used to process WES and RNA-seq data. FASTQ data were generated using the bcl2fastq tool (Illumina, v2.18) and then run through FASTQC for quality confirmation. For WES, reads were mapped to the human hg19 reference genome by the Burrows-Wheeler Alignment tool. The resulting bam files were processed using GATK best practice workflow (*25*). Copy number data was inferred from WES data through use of the CNVKit algorithm, using a pool of normal HapMap cell line samples as reference (*23*). The RSEM pipeline using STAR aligner was implemented to process RNA-seq data to get gene expression data (*24*).

In current cohort, fifty-eight PDX head and neck models were performed by WES and RNA-seq bioinformatics analysis. In each PDX model, it includes multiple (4 ≥ PDX) samples. For copy number data, consensus copy number status (2 = diploid, > 2 and < 5 = gain, and ≥ 5 = amplification) was called using majority voting among multiple PDX samples from same model. For gene expression data, average of Fragments Per Kilobase Million (FPKM) was taken to get gene expression at model level.

### Cell Lines

CAL33, BICR56, Detroit562, MSK921, BICR22, BEAS2B, 293T, and 293FT cell lines were obtained and maintained as previously described (*1*). All cells were incubated at 37 °C and 5% CO_2_. All cell lines were used in experiments for fewer than 20 passages (10 weeks) after thawing, before a fresh vial was taken from freeze. Cell lines in use were confirmed to be mycoplasma-negative using a Visual-PCR Mycoplasma Detection Kit (GM Biosciences).

### Establishment of Doxycycline-Inducible Knockdown Cell Lines

CAL33 and BICR56 inducible knockdown cells were generated by SIRION Biotech. MSK921 was generated in-house using lentiviral particles provided by SIRION (generated by transfection of 293TN cells with expression vectors and lentiviral packaging plasmids). The generation of the doxycycline-inducible knockdown cell lines were previously described (*1*). Doxycycline (Sigma-Aldrich) was used at 1 μg/mL to induce LZK knockdown.

### Generation of Tetracycline-Inducible Expression Cell Lines

The ViraPower HiPerform T-REx Gateway Expression System (Invitrogen) was used to generate cells with tetracycline-inducible expression of LZK WT and drug-resistant mutant (Q240S) as noted previously (*1*). Doxycycline (Sigma-Aldrich) was used at 1 μg/mL to induce LZK expression.

### Plasmids and Transfections

The stable overexpression of LZK using the pLenti6.3/TO/V5-DEST vector was described previously (*1*). The LZK Q240S drug-resistant construct and K195M kinase-dead mutant (*1*) was introduced using a Site-Directed Mutagenesis Kit (Stratagene) (Primers listed in Supplementary table S2). The LZK truncation constructs were amplified by PCR and then inserted into the pcDNA3.1-FLAG-HA vector using *XbaI* and *NotI* restriction sites (Primers listed in Supplementary table S3). 293T cells were transiently transfected using Lipofectamine 2000 (Invitrogen), according to the manufacturer’s protocol, with OptiMEM (Gibco). A pcDNA3.1-FLAG-HA vector (Addgene #52535) was used as an empty vector control where required. The CDK2 sensor vector CSII-pEF1a-DHB(aa994-1087)-mVenus and the nuclear marker vector CSII-pEF1a-H2B-mTurquoise were described previously (*22*).

### RNA Preparation

Cells were lysed using Buffer RLT (Qiagen) with 1% v/v 2-mercaptoethanol (Bio-Rad) 48 hours after treatment (tetracycline-induced overexpression or doxycycline-induced knockdown). Genomic DNA was removed, and RNA was prepared using a RNeasy kit (Qiagen) according to the manufacturer’s protocol. The RNA quantity was determined using a NanoDrop™ One Spectrophotometer (Thermo Scientific).

### RT-PCR

RT-PCR was performed using a SuperScript III One-Step RT-PCR kit (Invitrogen). Primers used are listed in Supplementary table S3. The cycling conditions for PCR were as follows: cDNA synthesis and pre-denaturation (one cycle at 55 °C for 30 minutes followed by 94 °C for two minutes), PCR amplification (25 cycles of denaturing at 94 °C for 15 seconds, annealing at 55 °C for 30 seconds, and extension at 68 °C for 60 seconds), and a final extension at 68 °C for five minutes using C1000 TOUCH CYCLER w/48W FS RM (Bio-Rad). PCR products were resolved on 2% agarose gel and visualized with Nancy-520 (Sigma-Aldrich) DNA gel stain under ultraviolet light using ChemiDoc™ MP Imaging System (Bio-Rad).

### Inhibitor Treatment

GNE-3511 was purchased from Cayman Chemical for *in vitro analyses* and Synnovator in large quantities for the *in vivo* mouse studies. MG132 was purchased from Selleck Chemicals. MLN4924 was purchased from MedChemExpress. All compounds were dissolved in DMSO (Fisher), and DMSO was used as the vehicle control in the cell-based assays.

### Protein Lysate Preparation, Immunoprecipitation, and Immunoblot Analysis

Generally, cells were plated in six-well or 35-mm plates for 24 hours, after which doxycycline was added for 48 hours or treatment with specific inhibitor was administered using 5% FBS media for specific time points. After appropriate treatment time, cells were washed with ice-cold phosphate-buffered saline without Ca and Mg (Quality Biological) and then lysed on ice for 10 minutes with RIPA buffer (50 mM NaCl, 1.0% IGEPAL^®^ CA-630, 0.5% sodium deoxycholate, 0.1% SDS, 50 mM Tris, pH 8.0) (Sigma-Aldrich) supplemented with protease inhibitor tablet (Sigma-Aldrich) and phosphatase inhibitor cocktails 2 and 3 (Sigma-Aldrich) followed by centrifugation at 15,000 rpm for 10 minutes at 4 °C. Protein concentrations were determined from the cell lysate by using 660 nm Protein Assay Reagent (Pierce). Cell extracts were denatured, subjected to SDS-PAGE, transferred to PVDF membranes (Bio-Rad), and blocked for 2 hours using 5% bovine serum albumin (BSA) in phosphate-buffered saline and 0.1% Tween® 20 (PBS-T). The membranes were incubated with the specific antibodies (as listed in Supplementary table S4) overnight in 5% BSA/PBST at 4 °C followed by 1 hour incubation with the appropriate horseradish peroxidase-conjugated secondary antibodies (Supplementary table S4) and signal was detected by chemiluminescence (Thermo Fisher).

For immunoprecipitation, cells were lysed in with 1x Triton X-100 cell lysis buffer (Cell Signaling Technology) supplemented with protease inhibitor tablet (Sigma-Aldrich) and phosphatase inhibitor cocktails 2 and 3 (Sigma-Aldrich). Cell lysates were calculated to 1 mg/mL followed by incubation with anti-FLAG (rat) antibody (BioLegend #637302) at 1:100 for two hours rotating at 4 °C. After, Dynabeads® Protein G (Life Technologies) was added for 1 hour rotating at 4 °C before washing four times with lysis buffer. Samples were denatured and immunoblot analysis was performed together with input samples at equal concentration.

### Subcellular Protein Fractionation

Proteins from the different cellular fractions, including cytoplasmic (CE), membrane (ME), soluble nuclear (SN), and chromatin-bound (CB) proteins, were isolated using Subcellular Protein Fractionation Kit For Cultured Cells (ThermoFisher Scientific) according to the manufacturer’s protocol. Briefly, cells were seeded, and the following day treated with doxycycline where appropriate, and incubated at 37 °C for 48 hours. Cellular compartments were sequentially extracted by incubating cells with cytoplasmic extraction buffer (CEB), followed by membrane extraction buffer (MEB), nuclear extraction buffer (NEB), and micrococcal nuclease containing NEB. Protein concentration was measured using 660 nm Protein Assay Reagent (Pierce) and immunoblot analysis was performed as described above.

### Reverse Phase Protein Arrays

Cells were seeded in 10 cm dishes, at 6 x 10^5^ for CAL33 and BICR56, and 6.25 x 10^5^ for MSK921, before addition of doxycycline (to induce LZK knockdown) the following day. Cells were lysed on ice with 1x Triton X-100 cell lysis buffer (Cell Signaling Technology) supplemented with protease and phosphatase inhibitors (Roche Applied Science) and 1.5 mM MgCl_2_, 48 hours after induction with doxycycline. Cell lysates were centrifuged, and the supernatant was collected. Protein concentration was measured using 660 nm Protein Assay Reagent (Pierce) and adjusted to 2 mg/mL. Then 4× reducing sodium dodecyl sulfate (SDS) sample buffer was added (40% glycerol, 8% SDS, and 0.25 M Tris HCl, pH 6.8, with 10% β-mercaptoethanol added before use), and the samples were incubated at 80 ^°^C for three minutes. Lysates from three independent experiments were sent for RPPA analysis. The Host and Tumour Profiling Unit at Cancer Research UK Edinburgh Centre (MRC Institute of Genetics and Molecular Mechanism, The University of Edinburgh) performed a nitrocellulose slide format RPPA with a panel of 60 antibodies according to established protocols (*40*). Results were compared to samples without dox-induction of LZK knockdown.

### Mass spectrometric analysis

Cells were seeded on a six-well plate at a density of 150,000 cells/well. The next day, cells were treated with doxycycline for 24 h, followed by treatment with 1 µM PROTAC97 or DMSO as a control. After 24 h, cells were lysed in lysis buffer (Cell Signaling Technology, catalog no. 9803S) and subjected to sample preparation and MS analysis.

Samples were subjected to chloroform/methanol precipitation, and the resulting protein pellets were washed twice with methanol. The protein pellets were air-dried for ∼10 min and resuspended in 100 mM HEPES (pH 8) by vigorous vortexing and sonication. Proteins were then reduced and alkylated with 10 mM tris(2-carboxyethyl)phosphine hydrochloride (TCEP)/40 mM chloroacetic acid (CAA) and digested with trypsin in a 1:50 (w/w) enzyme-to-protein ratio at 37°C overnight. Digestion was terminated by the addition of trifluoroacetic acid (TFA) to a 1% final concentration. The resulting peptides were labeled using an on-column TMT labeling protocol. TMT-labeled samples were compiled into a single TMT sample and concentrated using a SpeedVac concentrator. Peptides in the compiled sample were separated into eight fractions by off-line basic reversed-phase using the Pierce High pH Reversed-Phase Peptide Fractionation Kit (Thermo Fisher Scientific).

Prior to the liquid chromatography (LC)-MS measurement, the peptide fractions were resuspended in 0.1% TFA and 2% acetonitrile in water. Chromatographic separation was performed on an Easy-Spray Acclaim PepMap column (50 cm length × 75 µm inner diameter; Thermo Fisher Scientific) at 55°C by applying 120 min acetonitrile gradients in 0.1% aqueous formic acid at a flow rate of 300 nl/min. An UltiMate 3000 nano-LC system was coupled to a Q Exactive HF-X mass spectrometer via an easy-spray source (all Thermo Fisher Scientific). The Q Exactive HF-X was operated in TMT mode with survey scans acquired at a resolution of 60,000 at m/z 200. Up to 15 of the most abundant isotope patterns with charges 2-5 from the survey scan were selected with an isolation window of 0.7 m/z and fragmented by higher-energy collision dissociation with normalized collision energies of 32, while the dynamic exclusion was set to 35 s. The maximum ion injection times for the survey scan and dual MS (MS/MS) scans (acquired with a resolution of 45,000 at m/z 200) were 50 and 120 ms, respectively. The ion target value for MS was set to 3e6 and for MS/MS was set to 1e5, and the minimum AGC target was set to 1e3.

The data were processed with MaxQuant v. 1.6.17.0 (*41*) and the peptides were identified from the MS/MS spectra searched against Uniprot human reference proteome (UP000005640) using the built-in Andromeda search engine. Reporter ion MS2-based quantification was applied with reporter mass tolerance = 0.003 Da and min. reporter PIF = 0.75. Cysteine carbamidomethylation was set as a fixed modification and methionine oxidation, glutamine/asparagine deamination, as well as protein N-terminal acetylation, were set as variable modifications. For in silico digests of the reference proteome, cleavages of arginine or lysine followed by any amino acid were allowed (trypsin/P), and up to two missed cleavages were allowed. The false discovery rate (FDR) was set to 0.01 for peptides, proteins, and sites. A match between runs was enabled. Other parameters were used as pre-set in the software. Unique and razor peptides were used for quantification enabling protein grouping (razor peptides are the peptides uniquely assigned to protein groups and not to individual proteins). Reporter intensity corrected values for protein groups were loaded into Perseus v. 1.6.10.0 (*42*). Standard filtering steps were applied to clean up each dataset: reverse (matched to decoy database), only identified by site, and potential contaminant (from a list of commonly occurring contaminants included in MaxQuant) protein groups were removed. Reporter intensity corrected values were Log2 transformed. Protein groups with all values and at least 2 unique+razor peptides were kept. Reporter intensity values were then normalized by median subtraction within TMT channels. Student’s t-tests were performed on these values for groups of samples constituting a given dataset. The abundance changes thresholds of Log2 fold change (Log2FC) ≥ |1.0| and the significance threshold of –Log10 p-value ≥ 2.0 were applied to deliver protein groups with levels deemed reproducibly decreased or increased between the sample groups investigated.

### In Vitro Kinase Assay

One hundred nanograms of glutathione S-transferase (GST)-tagged human LZK pure protein (Carna Biosciences) was incubated with 100 ng of GST-tagged human inactive MKK7 pure protein (Carna Biosciences) in kinase buffer (Cell Signaling Technology). The assay was performed with 100 μM ATP at 37 °C for 30 minutes. Following the addition of 4× reducing SDS sample buffer, proteins were resolved by sodium dodecyl sulfate–polyacrylamide gel electrophoresis (SDS-PAGE) and immunoblot analysis was performed as stated previously.

### ELISA Assay

A PathScan® Phospho-SAPK/JNK (Thr183/Tyr185) Sandwich ELISA Assay (Cell Signaling Technology) was used for ELISA assays following the manufacturer’s protocol. In general, 500,000 cells were plated and treated with doxycycline the following day where appropriate and incubated at 37 °C for 48 hours. Cells were treated with the drug compound or control in 5% FBS media for 1 hour. After appropriate treatment time, cells were lysed on ice with 1x Cell Lysis Buffer (Cell Signaling Technology) supplemented with phosphatase and protease inhibitors (Sigma). Each diluted cell lysate was added to Phospho-SAPK/JNK (Thr183/Tyr185) Rabbit mAb Coated microwells in triplicate and incubated overnight at 4 °C. Samples were treated with the following antibodies and incubated at 37 °C for 1 hour and 30 minutes, respectively: Detection Antibody and HRP-Linked secondary antibody. Samples were washed between treatments using 1x Wash Buffer according to the manufacturer’s protocol. TMB substrate was added to each well and incubated at 37 °C for 10 minutes. Following this, STOP solution was added to each well and absorbance was measured at 450 nm using iMark™ Microplate Absorbance Reader (Bio-Rad). Graphs display relative phospho-JNK levels.

Eurofin’s KdELECT KINOMEscan^TM^ profiling

The binding affinity of GNE-3511, compound#21, and PROTAC-21A towards LZK was determined using Eurofin’s KdELECT KINOMEscan^TM^ assay. Briefly, kinase-tagged T7 phage strains were prepared in an E. coli host derived from the BL21 strain. The lysates were centrifuged and filtered to remove cell debris. The remaining kinases were produced in HEK-293 cells and subsequently tagged with DNA for qPCR detection. Streptavidin-coated magnetic beads were treated with biotinylated small molecule ligands for 30 minutes at room temperature to generate affinity resins for kinase assays. The liganded beads were blocked with excess biotin and washed with blocking buffer (SeaBlock (Pierce), 1% BSA, 0.05% Tween 20, 1 mM DTT) to remove unbound ligand and to reduce non-specific binding. Binding reactions were assembled by combining kinases, liganded affinity beads, and test compounds in 1x binding buffer (20% SeaBlock, 0.17x PBS, 0.05% Tween 20, 6 mM DTT). Test compounds were prepared as 111X stocks in 100% DMSO. Kds were determined using an 11-point 3-fold compound dilution series with three DMSO control points. The assay plates were incubated at room temperature with shaking for 1 hour and the affinity beads were washed with wash buffer (1x PBS, 0.05% Tween 20). The beads were then re-suspended in elution buffer (1x PBS, 0.05% Tween 20, 0.5 μM nonbiotinylated affinity ligand) and incubated at room temperature with shaking for 30 minutes. The kinase concentration in the eluates was measured by qPCR.

Eurofin’s scanMAX KINOMEscan^TM^ profiling

Thermodynamic interaction affinities between test compounds (compound#21 and PROTAC-21A) and >450 kinases were quantitatively measured using Eurofin’s scanMAX KINOMEscan^TM^ assay. Same experimental procedure was followed as the KdELECT profiling described above. Test compounds were prepared as 100x stocks in 100% DMSO and directly diluted into the assay. All reactions were performed in polypropylene 384-well plates in a final volume of 0.02 ml. The assay plates were incubated at room temperature with shaking for 1 hour and the affinity beads were washed with wash buffer (1x PBS, 0.05 % Tween 20). The beads were then re-suspended in elution buffer (1x PBS, 0.05 % Tween 20, 0.5 μM non-biotinylated affinity ligand) and incubated at room temperature with shaking for 30 minutes. The kinase concentration in the eluates was measured by qPCR.

### Immunofluorescence

Cells were seeded into each well of 8-well chamber slides and the following day treated with doxycycline where appropriate and incubated at 37 °C for 48 hours. Cells were then fixed in 4% paraformaldehyde for 5 min at 37°C and permeabilized with 0.3% Triton X-100 for 10 min at room temperature (RT). After blocking with 10% BSA + 0.1% Triton X-10 blocking solution for 30 min at RT, cells were co-incubated with primary rabbit anti-LZK antibody (1:50 dilution; Yen Zym) and mouse anti-p53 antibody (1:100 dilution; Santa Cruz Biotechnology) for 1 h, and secondary anti-rabbit Alexa Fluor549 and anti-mouse FITC-labeled antibodies (1:200 dilution; Invitrogen) for another hour at RT. The nuclei were stained with Hoechst 33342 (1:5000 dilution; Thermo Fisher Scientific) for 10 min at RT and the slides were observed using a Leico SP8 confocal microscope with a 63X oil immersion objective and appropriate filters. Images were analyzed using ImageJ/Fiji.

### Colony Formation Assays

Crystal violet assays were used to assess relative cell growth and survival after treatment with specific compounds. In general, cells were plated in triplicate in 12-well plates for 24 hours before drug treatments were added using 10% FBS media. The plates were incubated for 14 days, with the media and drug being replaced every 48 hours. For expression of the drug resistant LZK mutant (Q240S), doxycycline (0.25 μg/mL) was added and replaced every 48 hours with the drug or vehicle control. The cells were then washed with phosphate-buffered saline and fixed in ice-cold methanol before being stained with 0.5% crystal violet (Sigma-Aldrich) in 25% methanol. Images were taken using a ChemiDoc™ MP Imaging System (Bio-Rad), and for quantification, the crystal violet stain was dissolved in 33% acetic acid, incubated for 20 minutes with shaking, and read at 595 nm using iMark™ Microplate Absorbance Reader (Bio-Rad). Graphs display percent colony formation relative to the DMSO-treated control sample.

### Time-lapse Microscopy

Cells were plated in 96-well plates with full growth media more than 24 hours prior to imaging, such that the density would remain sub-confluent until the end of the imaging period. Time-lapse imaging was performed in 290 µL full growth media. Images were taken in CFP and YFP channels every 12 minutes on a Nikon Ti2-E inverted microscope (Nikon) with a 20X 0.45NA objective. Total light exposure time was kept under 600 milliseconds for each time point. Cells were imaged in a humidified, 37 °C chamber at 5% CO_2_.

### Image Analysis

All image analyses were performed with custom MATLAB scripts as previously described (*21*). In brief, optical illumination bias was empirically derived by sampling background areas across all wells in an imaging session and was subsequently used to flatten all images. This enabled measurement and subtraction of a global background for each image. Cells were segmented for their nuclei based on H2B-mTurquoise. CDK2 activity was calculated by measuring the nuclear and cytoplasmic fluorescence of the DHB-mVenus protein. Cells were segmented for their cytoplasmic regions by spatially approximating a ring with an inner radius of 2 µm outside of the nuclear mask and an outer radius with a maximum of 10 µm outside of the nuclear mask. Regions within 10 µm of another nucleus were excluded. Cytoplasmic DHB-mVenus was calculated as the median intensity within the cytoplasmic ring, excluding pixel intensities indistinguishable from background.

Mitosis events were automatically identified using H2B-mTurquoise and called at anaphase when one cell split into two daughter cells, each with approximately 45–55% of the size of the mother cell. Cells were considered to have been arrested in the G2 phase if a second mitosis was not detected for more than 30 hours after the first mitosis. Following drug treatment, cells were categorized by their CDK2 activity two hours after anaphase. Cells with high CDK2 activity (defined as greater than 0.6) are considered to have immediately re-entered the cell cycle, cells with transiently low CDK2 activity (defined as less than 0.6 two hours after anaphase but rising to greater than 0.6 within the viewing time) are considered to have entered into a transient G0 state before eventually re-entering the cell cycle, and cells with low CDK2 activity (defined as less than 0.6 within the viewing time) are considered to have entered a prolonged G0 state (*43*).

### EdU cell proliferation assay

Cell proliferation in response to GNE-3511 treatment was assessed by 5-ethynyl-2′-deoxyuridine (EdU) incorporation using the Click-iT EdU Alexa Fluor 488 cell proliferation assay kit (Invitrogen, Carlsbad, CA). CAL33 cells were seeded in µ-Plate 24 Well plates (Ibidi GmbH, Germany, Catalog # 82426) and treated with DMSO or GNE-3511 (100 nM, 250 nM, 500 nM, or 1 μM) for 48 hr after which the drug conditioned medium was replaced by 10 μM EdU medium for 30 min. Cell fixation, permeabilization, and EdU detection were performed according to the manufacturer’s instructions.

### Quantification and statistical analysis

All samples represent biological replicates. Data are presented as the mean with error bars shown on graphs representing ± SEM unless otherwise noted. Two-tailed Student’s *t*-test was used to assess significance of differences between groups for assays and used to measure significance of the mouse tumor volumes at the last day of treatment. Values of p < 0.05 were considered as significantly different.

### Chemical Synthesis

Reagents were purchased from commercial sources and used without further purification. *tert*-Butyl 4-(2-chloro-6-(3,3-difluoropyrrolidin-1-yl)pyridin-4-yl)piperidine-1-carboxylate was prepared as previously described(*18*). Synthetic details are provided for **PROTAC-21A**; PROTACs 21B-D were synthesized following a similar protocol. Final products were purified by flash chromatography or preparative high-performance liquid chromatography (HPLC). All tested compounds were characterized by liquid chromatography/mass spectrometry (LC/MS). Nuclear magnetic resonance (NMR) spectra were obtained on a 400 MHz Varian NMR and processed using MestreNova software (Mestralab). LC/MS data for small molecules were acquired on an Agilent Technologies 1290 Infinity HPLC system using a 6130 quadrupole LC/MS detector and a Poroshell 120 SB-C18 2.7 µm column (4.6 × 50 mm) or an Agilent 1200 series HPLC system with an LC/MSC Trap XCT detector and a ZORBAX 300SB-C18 3.5 µm column (4.6 × 50 mm). Preparative HPLC chromatography was performed on a Shimadzu system using a 30 mm × 150 mm Xbridge C18 column (Waters). Flash chromatography was performed on a Teledyne ISCO CombiFlash Rf+. High-resolution mass spectrometry (HRMS) data was acquired on a Waters Xevo G2-XS QTof running MassLynx version 4.1.

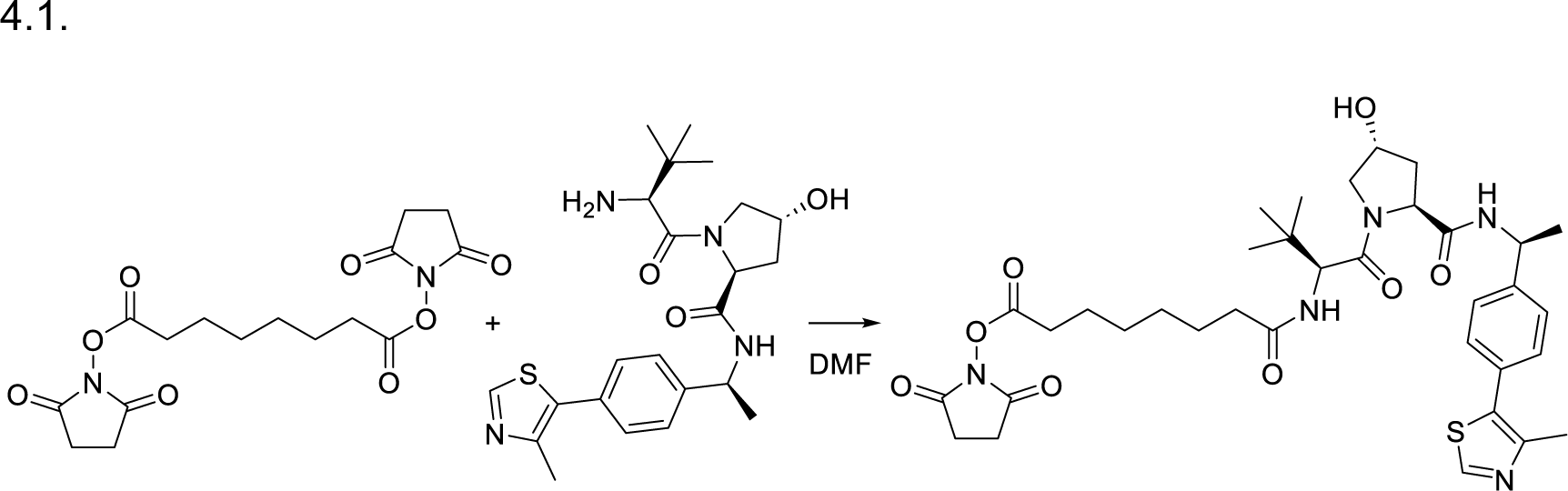

**2,5-Dioxopyrrolidin-1-yl 8-(((S)-1-((2S,4R)-4-hydroxy-2-(((S)-1-(4-(4-methylthiazol-5-yl)phenyl)ethyl)carbamoyl)pyrrolidin-1-yl)-3,3-dimethyl-1-oxobutan-2-yl)amino)-8-oxooctanoate.** A solution of 500 mg of (2S,4R)-1-((S)-2-amino-3,3-dimethylbutanoyl)-4-hydroxy-N-((S)-1-(4-(4-methylthiazol-5-yl)phenyl)ethyl)pyrrolidine-2-carboxamide hydrochloride salt and 0.22 mL of triethylamine in 2 mL of dimethylformamide (DMF) was added dropwise to a stirred solution of di-N-succinimidyl suberate (1.53 g, 4.16 mmol) in 10 mL of dry DMF and 10 mL of dry dichloromethane (DCM). The suspension was stirred overnight, then concentrated under reduced pressure. The residue was subjected to flash chromatography (0->15% MeOH in DCM) to yield 522 mg (72% yield) of the desired material as a colorless glassy residue. ^1^H NMR (400 MHz, CDCl_3_) δ 8.67 (br s, 1H), 7.43 (d, *J* = 7.8 Hz, 1H), 7.39 – 7.29 (m, 4H), 6.39 (d, *J* = 8.8 Hz, 1H), 5.04 (p, *J* = 7.0 Hz, 1H), 4.62 (t, *J* = 7.8 Hz, 1H), 4.56 (d, *J* = 8.9 Hz, 1H), 4.45 (tt, *J* = 4.9, 2.5 Hz, 1H), 4.18 (br s, 1H), 3.99 – 3.90 (m, 1H), 3.65 – 3.57 (m, 1H), 2.77 (s, 4H), 2.53 (t, *J* = 7.3 Hz, 2H), 2.47 (s, 3H), 2.33 (ddd, *J* = 12.7, 7.6, 4.7 Hz, 1H), 2.22 – 2.06 (m, 2H), 2.06 – 1.96 (m, 1H), 1.67 (p, *J* = 7.3 Hz, 2H), 1.57 (q, *J* = 7.3 Hz, 2H), 1.43 (d, *J* = 6.9 Hz, 3H), 1.40 – 1.20 (m, 4H), 0.99 (s, 9H).^13^C NMR (101 MHz, CDCl_3_) δ 173.41, 171.83, 170.12, 169.57, 168.69, 150.58, 148.28, 143.42, 131.71, 130.74, 129.53, 126.53, 69.87, 58.78, 57.43, 56.74, 48.78, 36.05, 35.95, 35.40, 30.85, 28.31, 28.09, 26.54, 25.67, 25.19, 24.42, 22.25, 16.05. HRMS: Calcd for C_35_H_48_N_5_O_8_S^+^ 698.3218, found 698.3215.

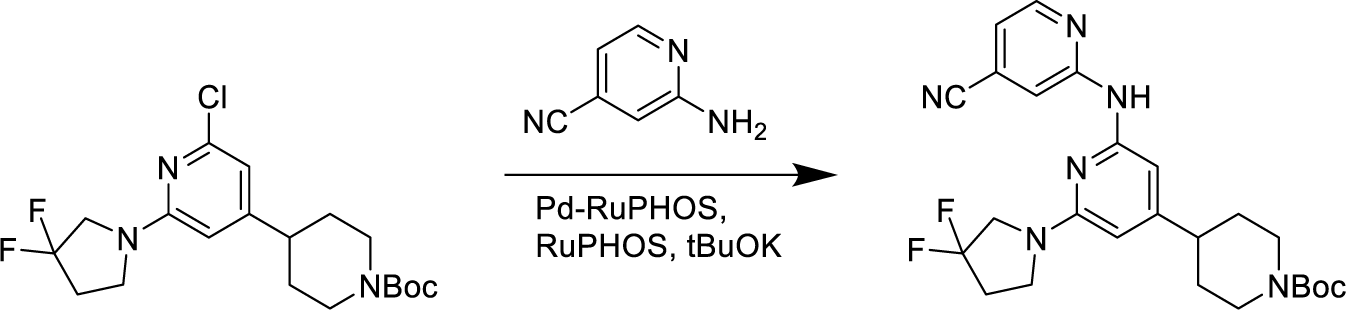

***tert*-Butyl 4-(2-((4-cyanopyridin-2-yl)amino)-6-(3,3-difluoropyrrolidin-1-yl)pyridin-4-yl)piperidine-1-carboxylate.** A microwave flask containing *tert*-butyl 4-(2-chloro-6-(3,3-difluoropyrrolidin-1-yl)pyridin-4-yl)piperidine-1-carboxylate (109 mg, 0.271 mmol), RuPhos (16.5 mg, 0.0353 mmol), chloro{[RuPhos][2-(2-aminoethylphenyl]palladium(II)}/[RuPhos] admixture (17.6 mg, 0.024 mmol), potassium t-butoxide (45.7 mg, 0.407 mmol), 2-amino-4-cyanopyridine (39.4 mg, 0.331 mmol), and a stir bar was sealed, and evacuated and backfilled with argon three times. Dioxane (3 mL) was added, and the mixture was heated for 45 minutes at 125 °C. Upon completion of the reaction, the mixture was adsorbed onto Celite and subjected to flash chromatography (15->60% EtOAc in hexanes). The product was obtained as a yellow solid (107 mg, 82% yield). ^1^H NMR (400 MHz, CDCl_3_) δ 8.42 (d, *J* = 1.2 Hz, 1H), 8.36 – 8.30 (m, 1H), 7.49 (s, 1H), 7.00 (dd, *J* = 5.1, 1.4 Hz, 1H), 6.22 (s, 1H), 5.80 (d, *J* = 1.0 Hz, 1H), 4.25 (br s, 1H), 3.84 (t, *J* = 13.1 Hz, 2H), 3.74 (t, *J* = 7.2 Hz, 2H), 2.79 (t, *J* = 12.7 Hz, 2H), 2.61 – 2.44 (m, 4H), 1.81 (d, *J* = 13.0 Hz, 2H), 1.68 – 1.52 (m, 2H), 1.48 (s, 9H). ^13^C NMR (101 MHz, CDCl_3_) δ 158.51, 156.16, 154.97, 154.66, 152.41, 149.20, 127.90 (t, *J* = 248 Hz), 121.52, 117.44, 117.07, 114.18, 98.82, 98.78, 97.48, 79.82, 54.33 (t, *J* = 32 Hz), 44.72, 44.26, 42.87, 34.26 (t, *J* = 24 Hz), 28.67. HRMS: Calcd for C_25_H_31_F_2_N_16_O_2_^+^ 485.2471, found 485.2468.

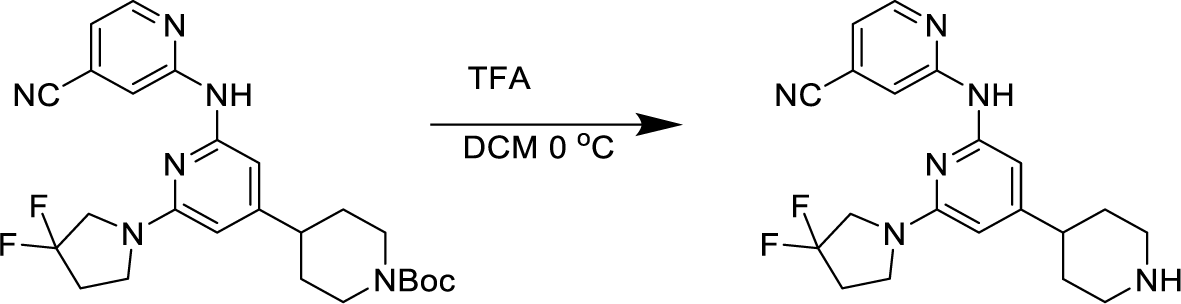

**2-((6-(3,3-Difluoropyrrolidin-1-yl)-4-(piperidin-4-yl)pyridin-2-yl)amino)isonicotinonitrile**. *tert*-Butyl 4-(2-((4-cyanopyridin-2-yl)amino)-6-(3,3-difluoropyrrolidin-1-yl)pyridin-4-yl)piperidine-1-carboxylate (81 mg, 0.167 mmol) was dissolved in 0.7 mL of DCM and stirred in an ice bath. Trifluoroacetic acid (0.7 mL) was added, and the reaction was stirred at 0 °C for 20 minutes. Upon completion of the reaction as determined by LC/MS, all volatiles were removed under reduced pressure. The resulting residue was used without further purification.

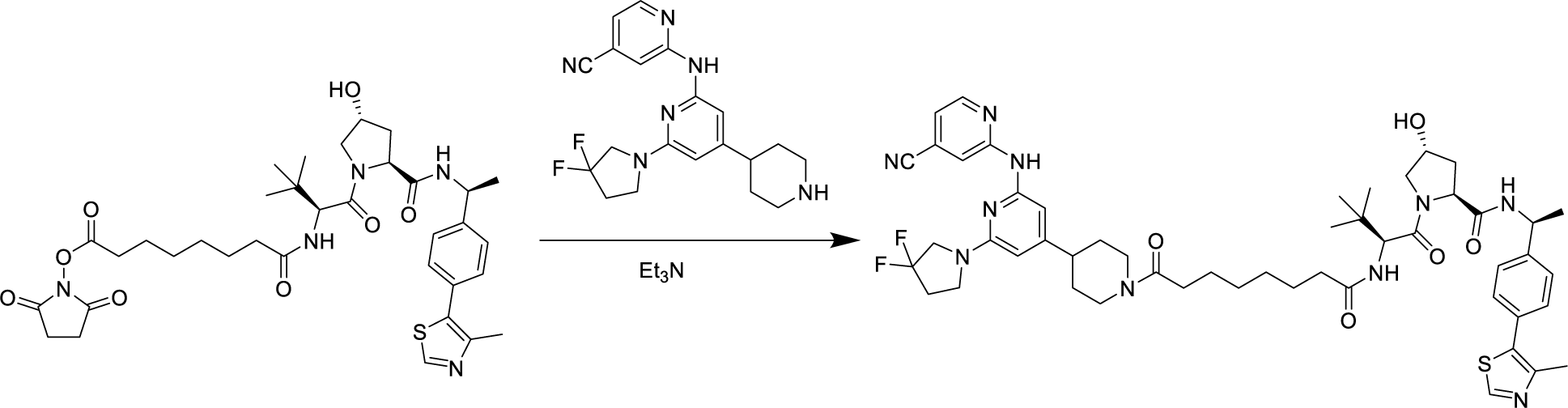

**PROTAC-21A.** 2,5-Dioxopyrrolidin-1-yl 8-(((S)-1-((2S,4R)-4-hydroxy-2-(((S)-1-(4-(4-methylthiazol-5-yl)phenyl)ethyl)carbamoyl)pyrrolidin-1-yl)-3,3-dimethyl-1-oxobutan-2-yl)amino)-8-oxooctanoate (522 mg, 0.75 mmol) was combined with 2-((6-(3,3-difluoropyrrolidin-1-yl)-4-(piperidin-4-yl)pyridin-2-yl)amino)isonicotinonitrile trifluoracetic acid salt (361 mg equivalent weight, 0.75 mmol) and N-methylmorpholine (0.5 mL) in 7 mL of DMF. After stirring overnight, the reaction was diluted with 75 mL each of DCM and water. The aqueous phase was separated and extracted with 2 × 50 mL DCM; the combined organic extracts were washed with saturated brine solution and dried over Na_2_SO_4_. Flash chromatography (5->25% MeOH in DCM) afforded 550 mg (76% yield) of the desired material as a yellow solid. ^1^H NMR (400 MHz, CDCl_3_) δ 8.65 (s, 1H), 8.46 (s, 1H), 8.28 (d, *J* = 5.1 Hz, 1H), 8.12 – 8.05 (br s, 1H), 7.57 (dd, *J* = 7.8, 3.9 Hz, 1H), 7.33 (m, 4H), 6.95 (d, *J* = 5.1 Hz, 1H), 6.61 (d, *J* = 8.9 Hz, 1H), 6.26 (s, 1H), 5.75 (s, 1H), 5.08 (p, *J* = 7.0 Hz, 1H), 4.72 (q, *J* = 7.8 Hz, 2H), 4.64 (d, *J* = 8.9 Hz, 1H), 4.48 (s, 1H), 4.29 (s, 1H), 4.02 (d, *J* = 11.3 Hz, 1H), 3.93 (d, *J* = 13.6 Hz, 1H), 3.80 (t, *J* = 13.0 Hz, 2H), 3.69 (t, *J* = 7.3 Hz, 2H), 3.62 (d, *J* = 10.7 Hz, 1H), 3.06 (t, *J* = 13.0 Hz, 1H), 2.65 – 2.35 (m, 6H), 2.32 (dt, *J* = 9.2, 4.3 Hz, 2H), 2.23 – 2.03 (m, 3H), 1.80 (d, *J* = 14.9 Hz, 2H), 1.63 – 1.41 (m, 10H), 1.29 (d, *J* = 7.2 Hz, 5H), 1.02 (s, 9H).^13^C NMR (101 MHz, CDCl_3_) δ 173.70, 172.09, 171.69, 170.14, 157.69, 155.95, 154.69, 152.59, 150.45, 148.92, 148.46, 143.41, 131.68, 130.82, 129.53, 127.80(t, *J* = 246 Hz), 126.54, 121.36, 117.33, 116.81, 114.34, 98.70, 97.31, 69.88, 58.87, 57.59, 56.93, 54.22(t, *J* = 32 Hz), 48.80, 46.16, 44.62, 42.76, 42.19, 36.32, 35.48, 34.11(t, *J* = 24 Hz), 33.27, 32.17, 29.03, 28.83, 26.66, 25.51, 25.20, 22.29, 16.17. HRMS: Calcd for C_51_H_65_F_2_N_10_O_5_S^+^ 967.4823, found 967.4814.

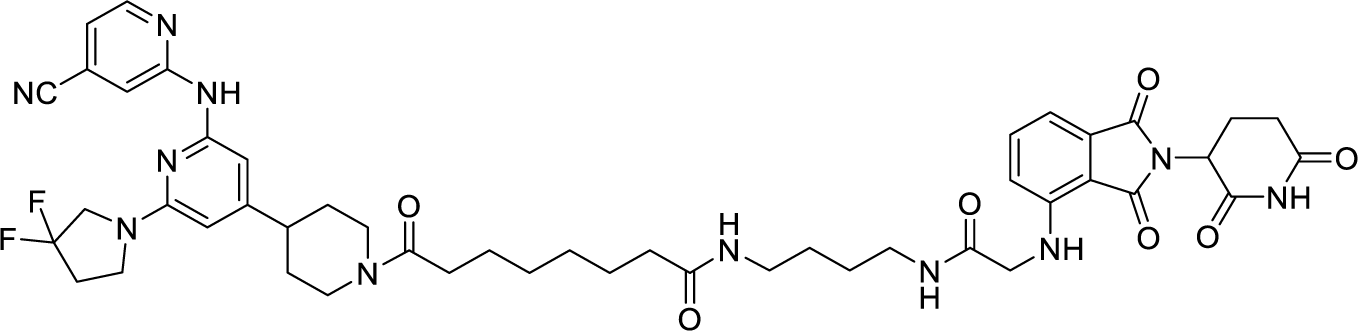

**PROTAC-21B: 98.4% pure (254 nm). HRMS: Calcd for C_47_H_56_F_2_N_11_O_7_^+^ 924.4332, found 924.4319.**

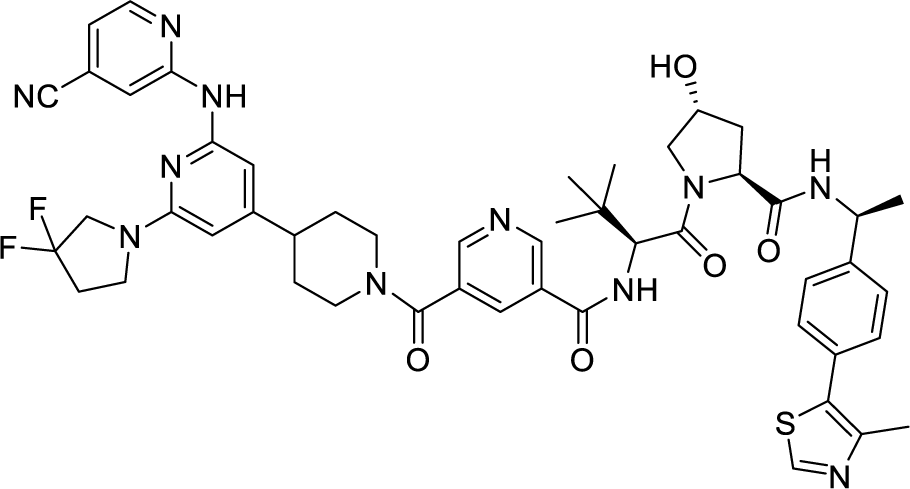

**PROTAC-21C: 97.4% pure (254 nm). HRMS: Calcd for C_50_H_56_F_2_N_11_O_5_S^+^ 960.4155, found 960.4153.**

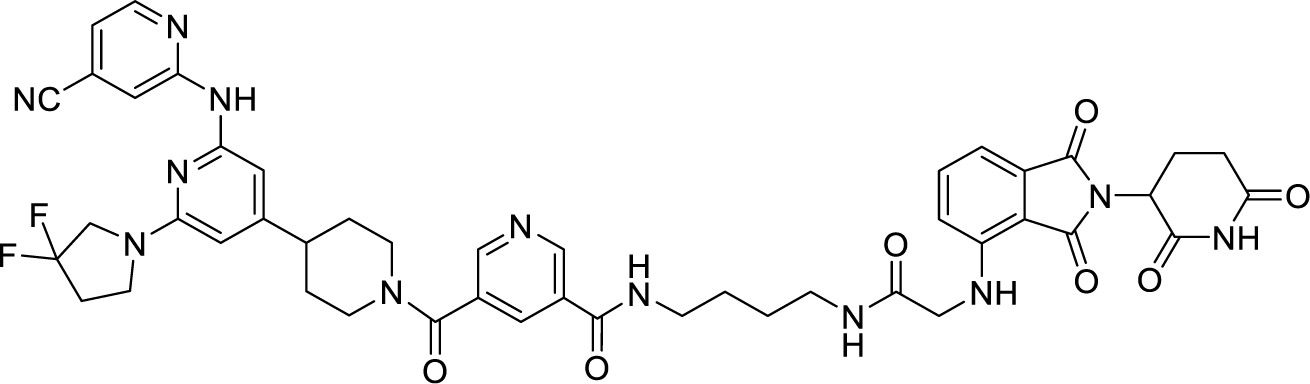

**PROTAC-21D: 95.4% pure (254 nm). HRMS: Calcd for C_46_H_47_F_2_N_12_O_7_^+^ 917.3659, found 917.3655.**

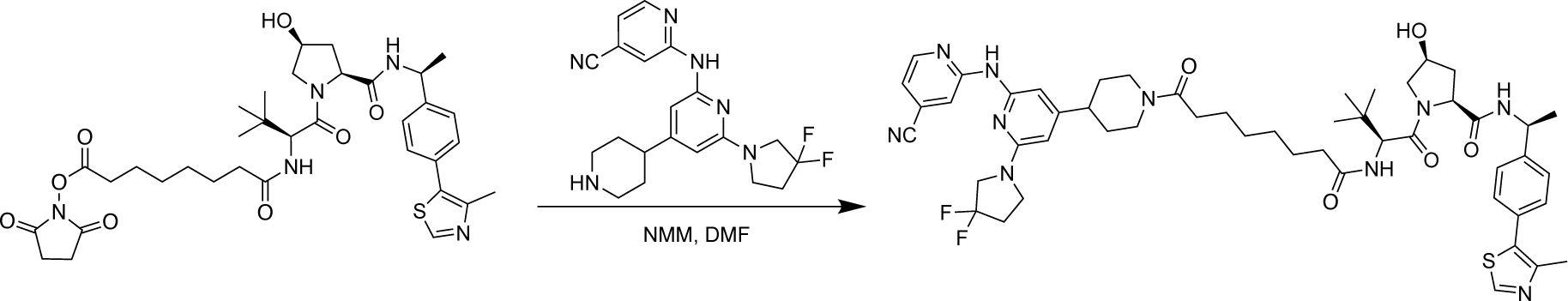

**PROTAC-21A cis-epimer**. The cis-epimer was prepared by similar method as described for VHL-suberoyl-DLK. To a solution of VHL-suberoyl-NHS (cis-epimer) (50 mg, 1 Eq, 72 μmol)in DMF (0.9 mL) was added 2-((6-(3,3-difluoropyrrolidin-1-yl)-4-(piperidin-4-yl)pyridin-2-yl)amino)isonicotinonitrile (28 mg, 1.0 Eq, 74 μmol) (TFA salt) and 4-methylmorpholine (43 mg, 47 μL, 6 Eq, 0.43 mmol). The reaction was stirred overnight and diluted with water and dichloromethane (20 mL). The aqueous phase was separated and further extracted twice with dichloromethane (2 x 20 mL). The combined extracts were washed with saturated brine solution, dried over sodium sulfate, filtered, and concentrated under reduced pressure. The residue was purified by reverse phase HPLC (Water/Acetonitrile (0.05%TFA)). Desired fractions were passed through a Stratosphere bicarbonate column (0.36 mmol), concentrated under reduced pressure and lyophilized to give the final compound as a yellow solid (39.8 mg, 72 μmol, 57% yield) ^1^H NMR (400 MHz, CDCl_3_) δ 8.66 (s, 1H), 8.42 (s, 1H), 8.31 (d, *J* = 5.1 Hz, 1H), 7.67 (d, *J* = 7.9 Hz, 1H), 7.60 (d, *J* = 8.7 Hz, 1H), 7.42 – 7.31 (m, 4H), 6.98 (dd, *J* = 5.1, 1.4 Hz, 1H), 6.21 (s, 1H), 6.15 (d, *J* = 9.1 Hz, 1H), 5.76 (s, 1H), 5.44 (d, *J* = 9.4 Hz, 1H), 5.07 (p, *J* = 7.0 Hz, 1H), 4.75 (dd, *J* = 15.9, 10.6 Hz, 2H), 4.59 (d, *J* = 9.1 Hz, 1H), 4.47 – 4.39 (m, 1H), 4.00 – 3.89 (m, 2H), 3.85 – 3.77 (m, 3H), 3.71 (t, *J* = 7.2 Hz, 2H), 3.09 (td, *J* = 13.2, 2.5 Hz, 1H), 2.68 – 2.42 (m, 7H), 2.38 – 2.26 (m, 3H), 2.24 – 2.18 (m, 2H), 2.13 (ddd, *J* = 14.1, 9.2, 5.0 Hz, 1H), 1.86 (t, *J* = 15.4 Hz, 2H), 1.63 (t, *J* = 7.3 Hz, 4H), 1.56 (dd, *J* = 12.7, 3.7 Hz, 1H), 1.48 (d, *J* = 6.9 Hz, 3H), 1.40 – 1.31 (m, 4H), 1.04 (s, 9H). ^13^C NMR (101 MHz, CDCl_3_) δ 173.20, 172.53, 171.66, 171.53, 157.66, 156.14, 154.58, 152.53, 150.48, 149.09, 148.68, 142.47, 131.53, 131.29, 130.26, 129.74, 127.80, 126.58, 125.34, 121.40, 117.35, 116.96, 114.14, 98.59, 97.29, 71.09, 60.01, 58.70, 57.07, 54.23 (t, *J* = 31.9 Hz), 49.33, 46.16, 44.62, 42.84, 42.17, 36.49, 35.22, 34.99, 34.15 (t, *J* = 23.9 Hz), 33.34, 33.28, 29.13, 29.01, 25.51, 25.28, 21.97, 16.19. HRMS: Calcd for C_51_H_65_F_2_N_10_O_5_S+ : 967.4828, found, 967.4826.

## Supplementary Materials

Fig S1. LZK inhibitor diminishes growth of HNSCC cell lines.

Fig S2. Kinase-dependent and - independent roles of LZK affect c-MYC and p53 abundance, respectively

Fig S3. Inhibition of LZK impairs progression through the cell cycle

Fig S4. Inhibition of LZK does not induce apoptosis in HNSCC cell lines

Fig S5. GNE-3511 at 50 mg/kg is well tolerated in the PDX mouse models

Fig S6. Compound #21 inhibits LZK activity and decreases viability of HNSCC cells

Fig S7. Dependence of PROTAC-21A-mediated degradation of LZK involves both ubiquitin-like molecule NEDD8 and the proteasome

Fig S8. TREEspot^TM^ interaction maps for compound #21 and PROTAC-21A

Fig S9. Shotgun proteomic analysis of total cell protein extracts from CAL33 TR LZK WT cell line following treatment with PROTAC97

Fig S10. LZK-targeting PROTAC reduces HNSCC viability but exhibits low membrane permeability Table S1. *MAP3K13* amplification status for the HNSCC PDX models used in this study.

Table S1. *MAP3K13* amplification status for the HNSCC PDX models used in this study.

Table S2. List of oligonucleotides used in this study.

Table S3. Primer sets for RT-PCR analyses.

Table S4. List of antibodies used for western blot analysis.

Data File S1. Kinomescan matrix for compound #21 and PROTAC-21A

Data File S2. Shotgun proteomic analysis of protein levels in the total cell extracts from LZK expressing CAL33 cells

## Acknowledgments

We thank the NCI Patient-Derived Models Repository (PDMR) for the PDX models (391396-364-R, 295223-140-R, 226611-082-R, 328373-195-R, and 959717-210-R) and providing the NGS and RNA-seq data for the fifty-eight HNSCC PDX mouse models for bioinformatic analysis. Also, we thank the members of the Brognard laboratory and the Laboratory of Cell and Developmental Signaling for all of the helpful advice.

## Author contributions

- Conception and design: A.L. Funk, M. Katerji, M. Afifi, S. D. Cappell, C.C. Woodroofe, R.E. Swenson, and J. Brognard
- Development of methodology: A.L. Funk, M. Katerji, M. Afifi, S. D. Cappell, C.C. Woodroofe, R.E. Swenson, and J. Brognard
- Acquisition of data: A.L. Funk, M. Katerji, K. Nyswaner, C.C. Woodroofe, Z.C. Edwards, M. Afifi, K.L. Bergman, E.W. Trotter, M. R. Rubin, A. Marusiak. K. Karpińska, E. Lindberg
- Analysis and interpretation of data: A.L. Funk, M. Katerji, K. Nyswaner, C.C. Woodroofe, M. Afifi, A. Marusiak. K. Karpińska, E. Lindberg, S.D. Cappell, R.E. Swenson, and J. Brognard
- Writing, review and/or revision of the manuscript: A.L. Funk, N. R. Gough, K. Nyswaner, C.C. Woodroofe, Z.C. Edwards, K.L. Bergman, M. Katerji, M. R. Rubin, I. Ahel, S. Dash, M.Afifi, S. Cappell, R.E. Swenson, and J. Brognard
- Study supervision: A.L. Funk, M. Katerji, and J. Brognard
- Other: Execution of the HNSCC PDX *in vivo* experiments with GNE-3511: A.L. Ries, A. James, C.M. Robinson, and S. Difilippantonio; Performed HNSCC PDX; NGS data analysis of PDMR HNSCC PDX mouse models: T.C. Chang, J.H. Doroshow, and L. Chen; Pharmacokinetic analysis of LZK PROTACs: X. Xu.

## Data and Materials Availability

Data from mass spectrometry are deposited in a publicly accessible repository and available via ProteomeXchange Consortium (*44*) via the PRIDE partner repository (*45*) with the dataset identifier PXD054481.

---

**Supplementary Figure S1.**
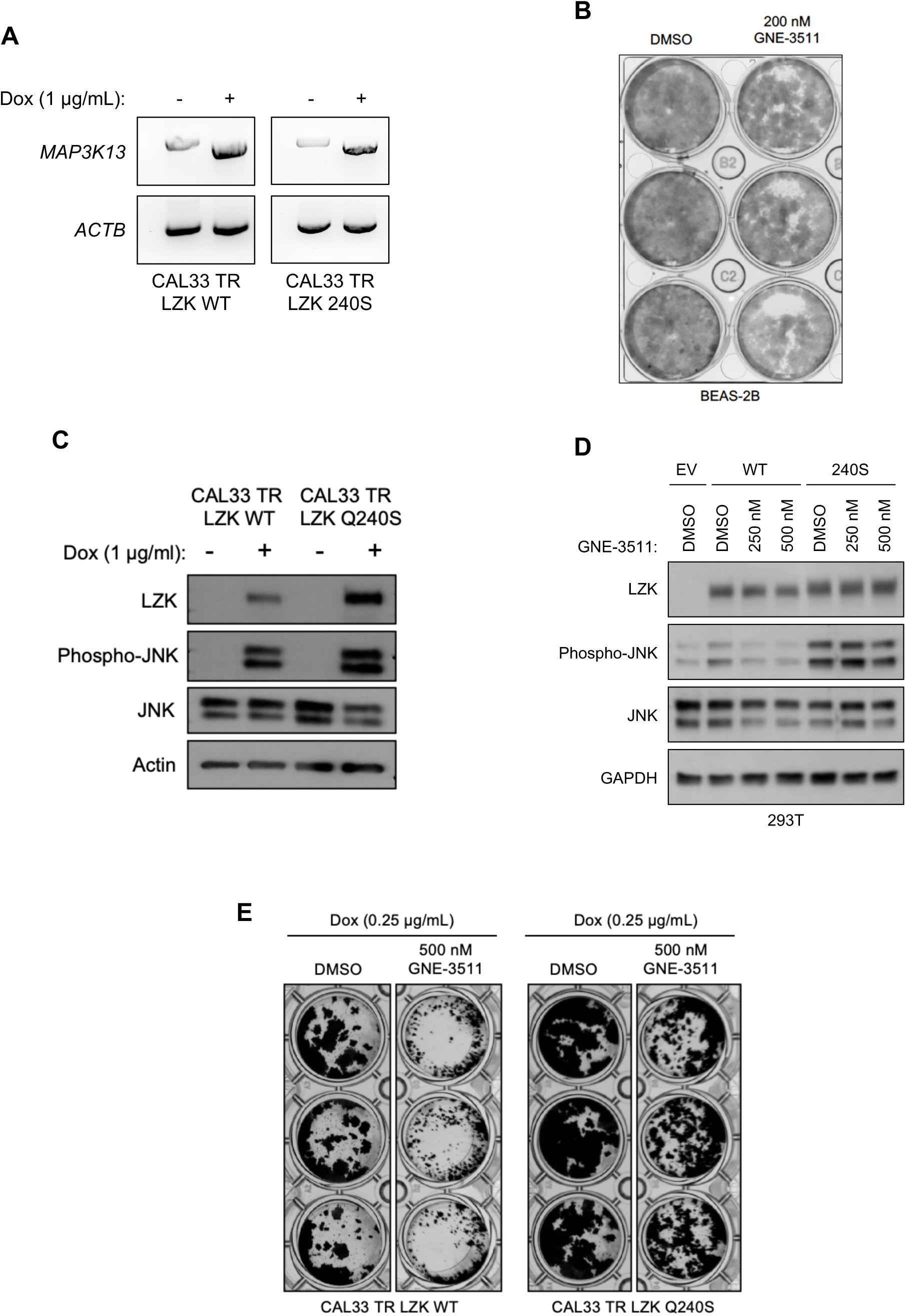
LZK inhibitor diminishes growth of HNSCC cell lines. **A.** RT-PCR analysis showing *MAP3K13* transcripts of dox-inducible wild-type (LZK WT) or drug-resistant form (LZK Q240S) in stable CAL33 cell lines. **B.** Effect of GNE-3511 on colony formation of normal bronchial epithelial cells BEAS-2B. Data are representative of 3 independent experiments. **C.** Western blot analysis showing LZK and pJNK expression levels in dox-inducible wild-type (LZK WT) or drug-resistant LZK (LZK Q240S) expressing CAL33 cell lines. Data are representative of 3 independent experiments. **D.** Western blot showing the LZK dependence of the inhibitory effect of GNE-3511 on phosphorylation of JNK in 293T cells expressing either LZK WT or LZK Q240S. GAPDH served as the loading control. Data are representative of 3 independent experiments. EV, Empty Vector. **E.** Effect of GNE-3511 on colony formation of CAL33 cells with dox-induced LZK WT vs Q240S. Data are representative of 3 independent experiments.

**Supplementary Figure S2.**
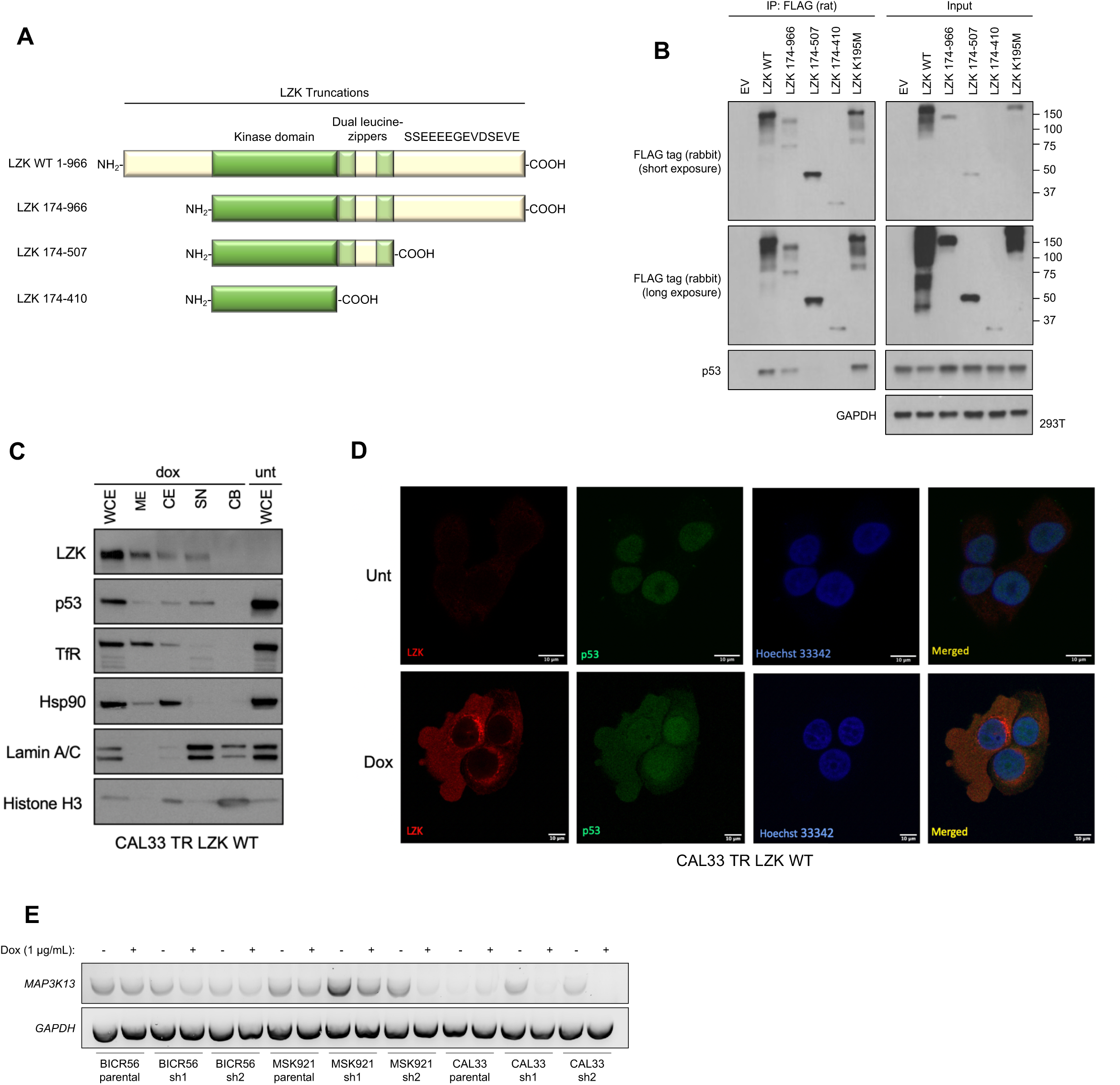
Kinase-dependent and - independent roles of LZK affect c-MYC and p53 abundance, respectively. **A.** Schematic diagram of the N-terminal FLAG-tagged LZK and its truncation constructs tested for interaction with p53. **B.** Western blot showing interaction between p53 and the indicated FLAG-tagged LZK constructs as determined by co-immunoprecipitation with a FLAG antibody (left). Long and short exposures of the Western blot for FLAG-tagged LZK are shown. Input shows the abundance of the indicated proteins in the lysate used for co-immunoprecipitation. Data are representative of 3 independent experiments. **C.** Western blot showing LZK and GOF-p53 in membrane (ME), cytoplasmic (CE), soluble nuclear (SN) and chromatin-bound nuclear (CB) extracts from CAL33 TR LZK WT cells. Transferrin Receptor (TfR) served as the marker of ME compartment, Hsp90 as the marker of CE compartment, Lamin A/C as the marker of SN compartment, and Histone H3 as the marker of CB compartment. Data are representative of 3 independent experiments. **D.** Images of dox-induced CAL33 TR LZK WT cells stained for GOF-p53 (green) and LZK (red). The nuclei were stained with Hoechst 33342. Merged image shows colocalization. Scale bar = 10 µM. Data are representative of 3 independent experiments, each assessing 10 individual cells. **E.** RT-PCR analysis showing *MAP3K13* transcripts in the indicated parental cell lines or cell lines with dox-inducible shRNA targeting LZK. Data are representative of 3 independent experiments.

**Supplementary Figure S3.**
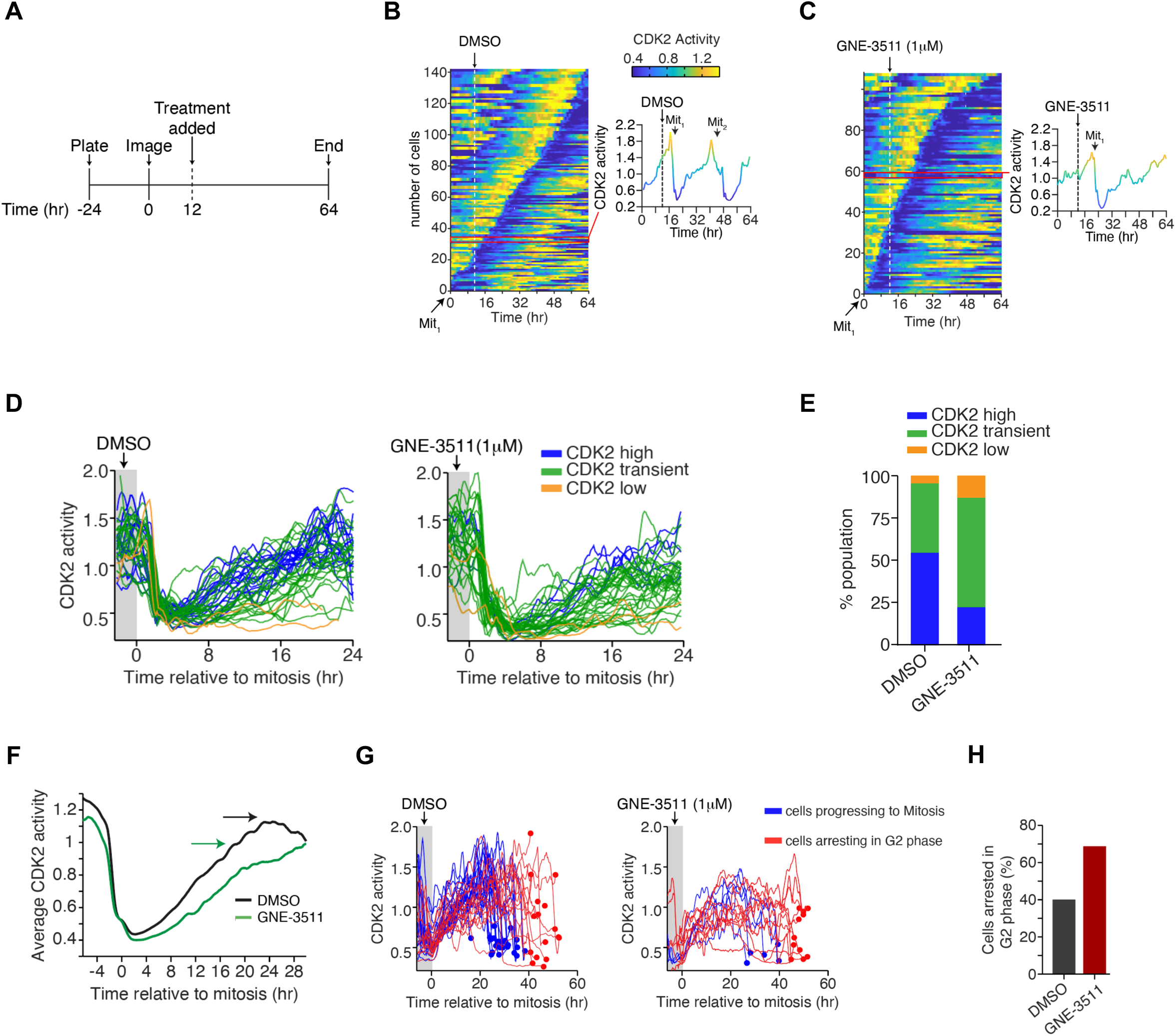
Inhibition of LZK impairs progression through the cell cycle. **A**. Diagram of experimental protocol in SCC15 cells. Mitotic events were monitored using the peaks of CDK2 activity in each cell. Images were acquired 12 hours before exposure to treatment (DMSO or GNE-3511) and then every 12 minutes for 52 more hours. **B, C.** Heat maps of CDK2 activity in asynchronously cycling cells treated with DMSO (vehicle control) (B) or GNE-3511 (C) at the indicated time. Cells are sorted by the time of the first mitosis relative to the start of the imaging. Although the time of mitosis was independently identified using the nuclear marker H2B-mTurquoise, it can also be visualized by a rapid drop in CDK2 activity, as indicated by the black arrow. Inset is a CDK2 activity trace for a single representative cell. The red line indicates the position of that cell within the heat map. Mitotic events are noted as Mit_1_ (first division after exposure), and Mit_2_ (second division after exposure). Data are representative of 2 independent experiments. **D, E.** Effect of DMSO (vehicle control) and GNE-3511 on CDK2 activity throughout the cell cycle. Panel E represents a quantitative bar graph showing the percentage of the cell populations relative to CDK2 activity in treated cells. 2 independent experiments. **F.** Graph showing the effect of DMSO or GNE-3511 on average CDK2 activity during progression through the cell cycle. Arrows indicate maximum CDK2 activity reached. **G, H.** Graph showing the effect of DMSO and GNE-3511 on CDK2 activity in cells progressing to mitosis or arresting in G2-phase. Panel H represents a quantitative bar graph showing the percentage of cells arrested in G2 phase. Data are representative of 2 independent experiments.

**Supplementary Figure S4.**
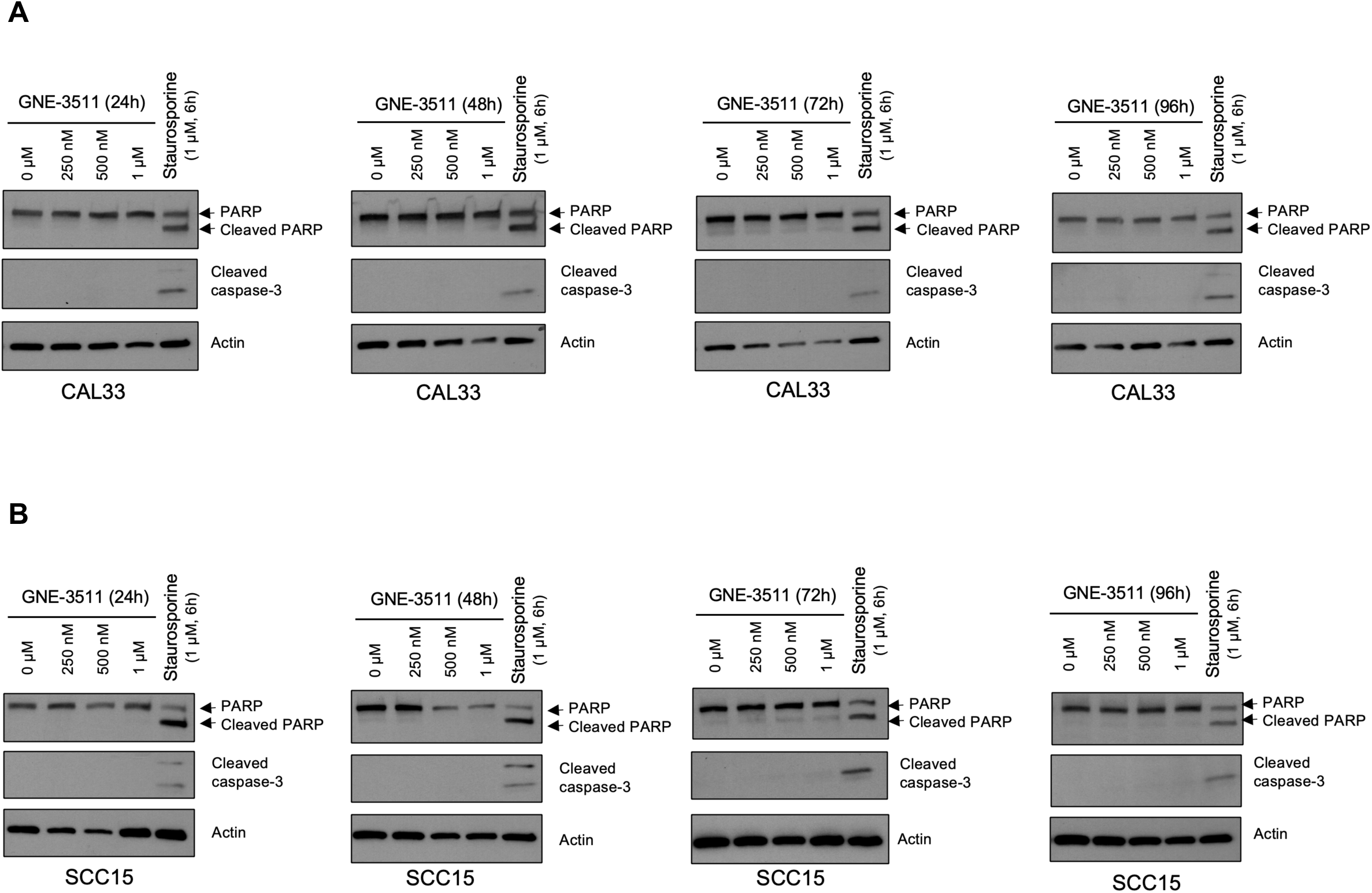
Inhibition of LZK does not induce apoptosis in HNSCC cell lines. A, **B.** Western blots showing the effect of increasing concentrations of GNE-3511 treatment on apoptotic markers cleaved PARP and cleaved caspase-3 in CAL33 (A) and SCC15 (B) cells at different time-points (24h-96h). Six-hour treatment of 1 µM Staurosporine was used as a positive control to induce apoptosis. Actin served as a loading control. Data are representative of 3 independent experiments.

**Supplementary Figure S5.**
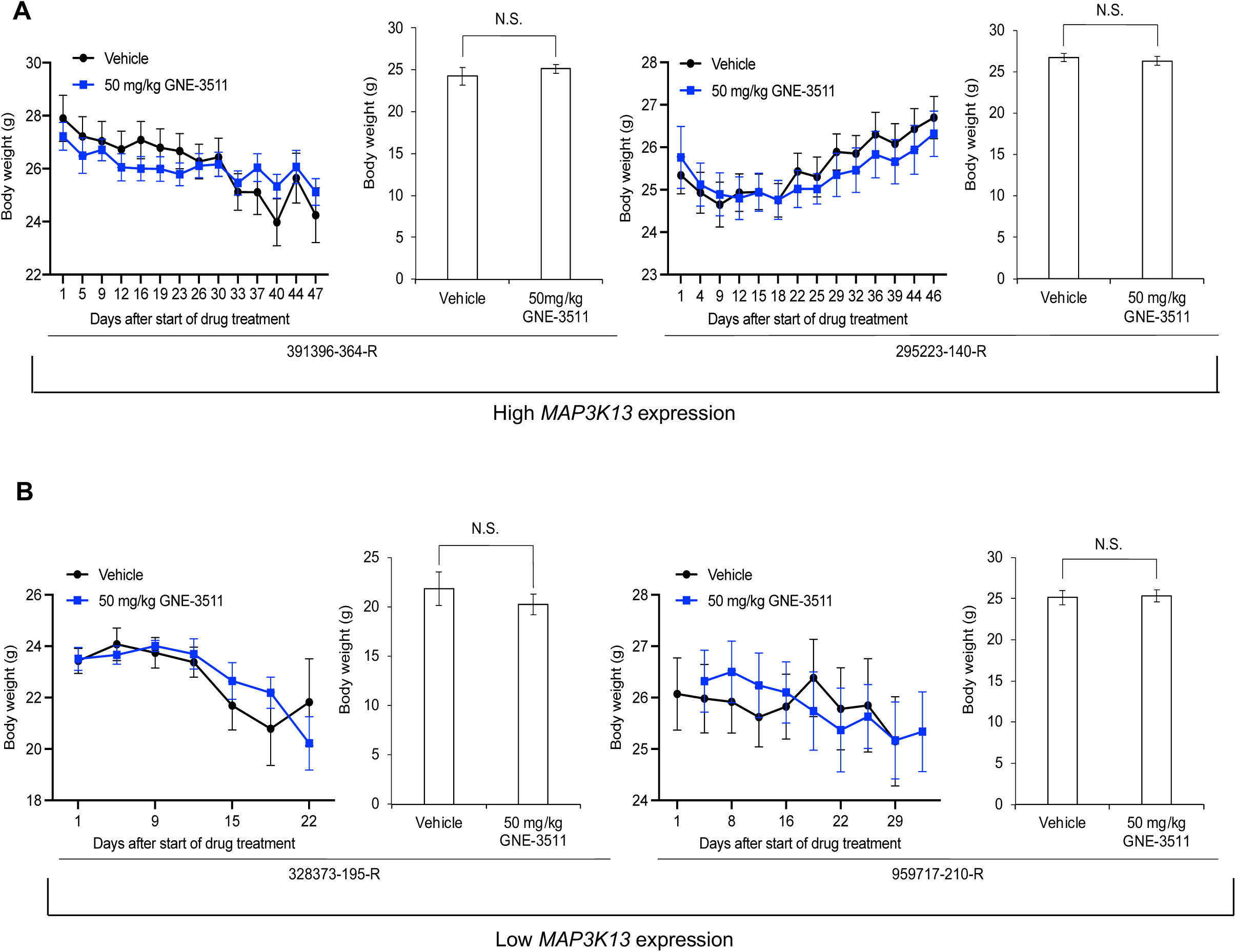
GNE-3511 at 50 mg/kg is well tolerated in the PDX mouse models. **A.** Body weights of the mice for the HNSCC PDX mouse models harboring *MAP3K13* amplification (PDX #: 391396-364-R and 295223-140-R) treated with vehicle or GNE-3511 (50 mg/kg, q.d., five days on/two days off). Mean mouse body weight ± SEM are shown; Student’s *t*-test; N.S., not significant. **B.** Body weights of the mice for the HNSCC PDX mouse models lacking amplification (PDX #: 328373-195-R and 959717-210-R) treated with vehicle or GNE-3511 (50 mg/kg, q.d., five days on/two days off). Mean mouse body weight ± SEM are shown; Student’s *t*-test; N.S., not significant.

**Supplementary Figure S6.**
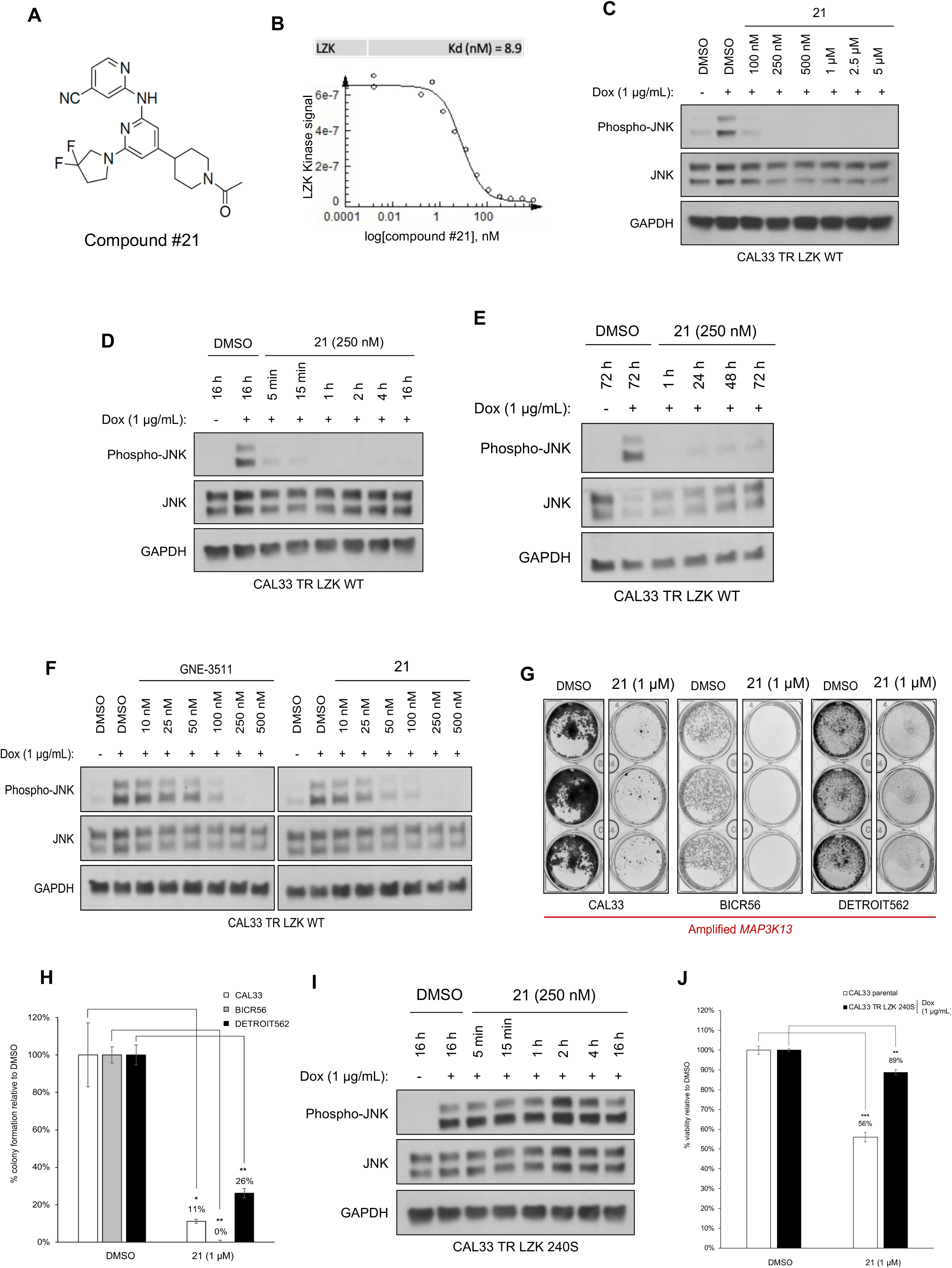
Compound #21 inhibits LZK activity and decreases viability of HNSCC cells. **A**. Chemical structure of compound #21, the pharmacophore used in LZK-targeting PROTACs. **B.** Dose curve of the *in vitro* binding affinity of compound #21 to LZK assessed by Eurofin’s KdELECT KINOMEscan^TM^ profiling. The amount of kinase measured by qPCR (Signal; y-axis) is plotted against the corresponding compound concentration in nM in log10 scale (x-axis). **C.** Western blot showing the effect of increasing concentrations of compound #21. LZK WT was induced with dox in CAL33 cells and the cells were exposed to the indicated concentration of #21 and phosphorylated JNK was monitored. GAPDH served as the loading control. Data are representative of 3 independent experiments. **D, E.** Western blots showing the effect of compound #21 over the indicated length of exposure in CAL33 LZK WT cells. Phosphorylated JNK was monitored. GAPDH served as the loading control. Data are representative of 3 independent experiments. **F.** Western blots showing the effect of GNE-3511 or compound #21 on phosphorylated JNK levels in dox-induced CAL33 LZK WT cells exposed to the indicated concentrations of drug. GAPDH served as the loading control. Data are representative of 3 independent experiments. **G, H.** Effect of compound #21 on colony formation of the indicated HNSCC cell lines. Panel G shows a representative experiment; panel H shows quantitative data for 3 experiments presented as the mean ± SEM; Student’s *t*-test; \*\**p* < 0.01, \**p* < 0.05. **I.** Western blot showing the effect of compound #21 on phosphorylated JNK levels in CAL33 LZK^Q240S^ drug-resistant mutant cells exposed for up to 16 hours. GAPDH served as the loading control. Data are representative of 3 independent experiments. **J.** Effect of compound #21 on cellular viability in CAL33 cells with or without overexpression of LZK Q240S. Expression of LZK Q240S was induced with dox. Viability was assessed 72 hours after addition of compound #21 or DMSO (as the vehicle control) and determined by MTS assay. Data are shown as mean ± SEM for 3 experiments with triplicates each. Student’s *t*-test; \*\*\**p* < 0.001, \*\**p* < 0.01.

**Supplementary Figure S7.**
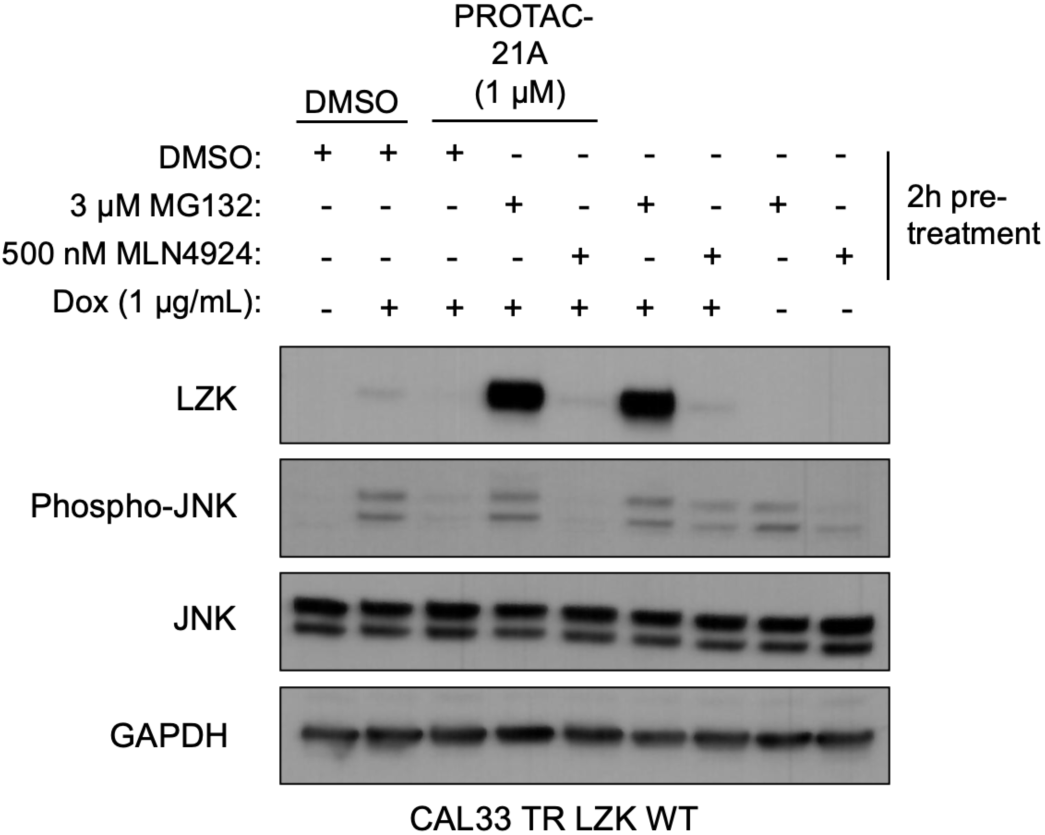
Dependence of PROTAC-21A-mediated degradation of LZK involves both ubiquitin-like molecule NEDD8 and the proteasome. LZK WT was induced with dox in CAL33 cells and the cells were pre-treated for 2 hours with either MG132 or MLN4924 alone, or in combination with PROTAC-21A treatment for 24 h. Expression levels of LZK and phosphorylated JNK were monitored. GAPDH served as the loading control. Data are representative of 3 independent experiments.

**Supplementary Figure S8.**
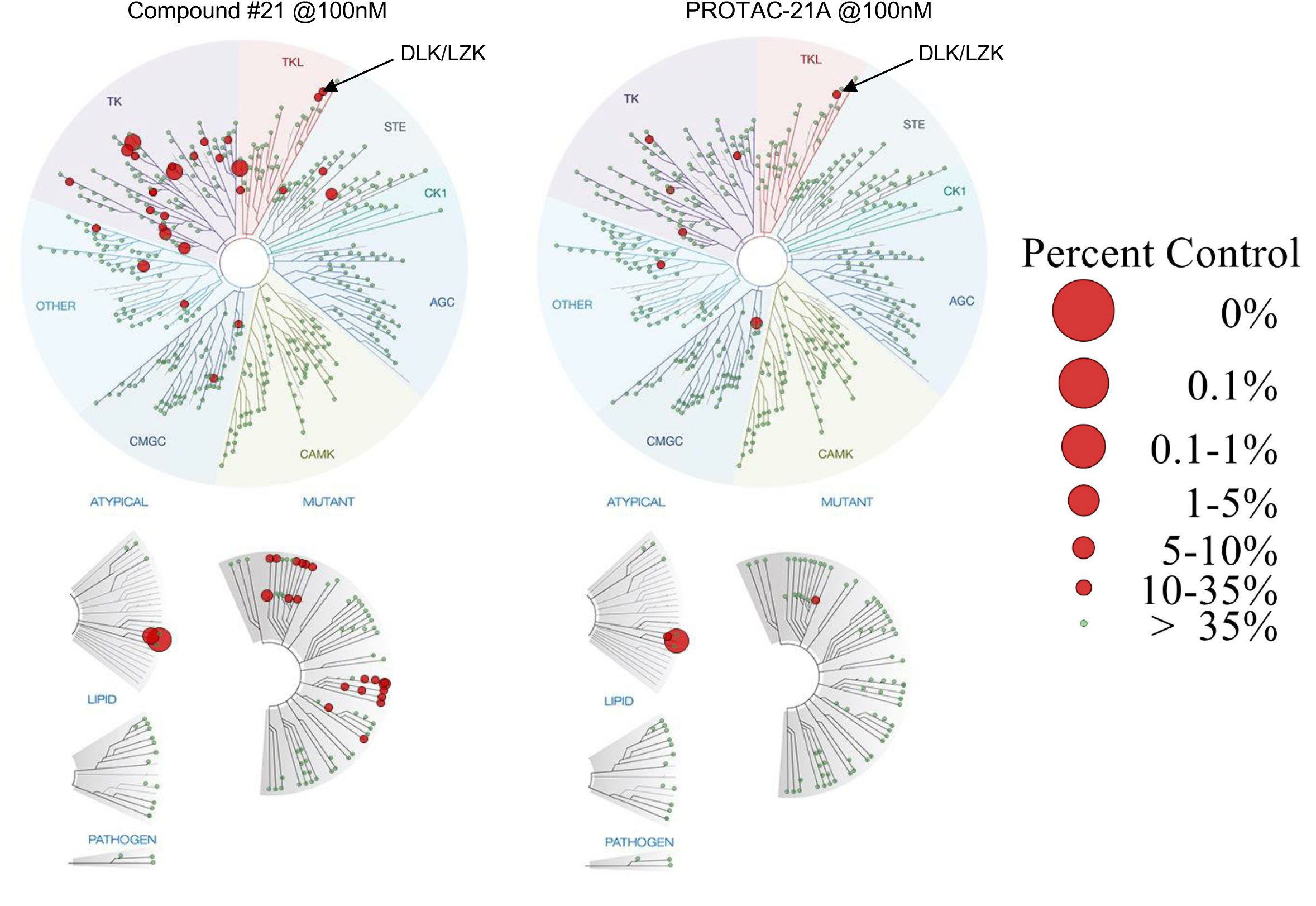
TREE*spot*^TM^ interaction maps for compound #21 and PROTAC-21A. KINOMEscan screening was performed for compound#21 and PROTAC-21A using an active site-directed ATP-independent competition binding assay by Eurofins discovery that quantitatively measures interactions between the two test compounds and more than 450 human kinases and disease relevant mutant variants. Both compounds were screened at 100nM, and results for primary screen binding interactions are reported as % Ctrl where lower numbers indicate stronger hits in the matrix. Kinases found to bind are marked with red circles, where larger circles indicate higher-affinity binding. Arrow indicates LZK/DLK binding. The kinomescan matrix for compound #21 and PROTAC-21A are represented in data file S1.

**Supplementary Figure S9.**
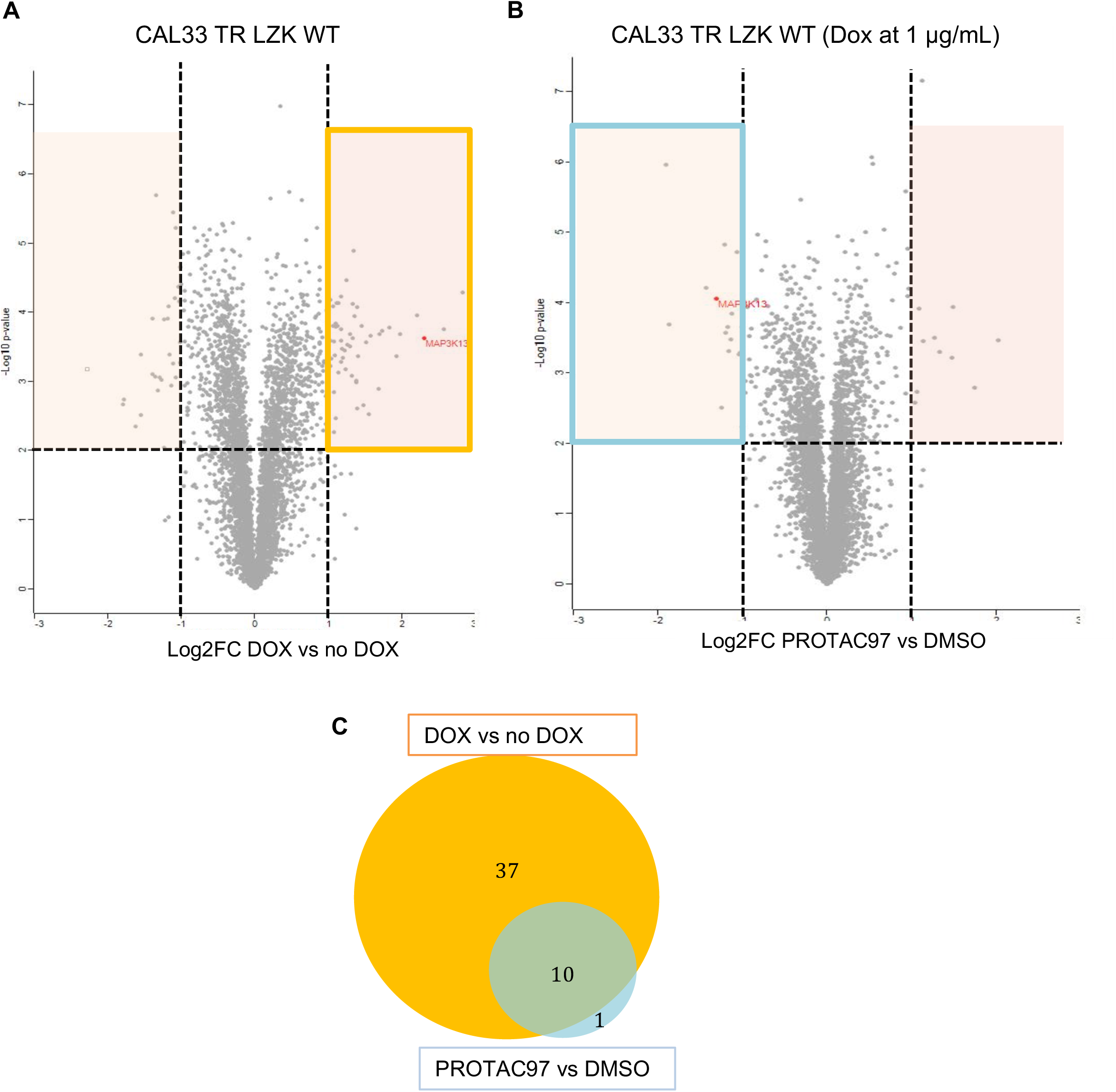
Shotgun proteomic analysis of total cell protein extracts from CAL33 TR LZK WT cell line following treatment with PROTAC97. **A, B.** CAL33 TR LZK WT cells were treated with doxycyline (DOX) to induce LZK overexpression, followed by treatment with 1 µM PROTAC97 or DMSO control for 24 h. LZK protein is indicated in red. Volcano plot displaying the log2 fold change (Log2FC, x axis) against the t-test-derived −log10 statistical (n=3) p-value (y axis) for all proteins detected in the total cell extract from CAL33 TR LZK WT cells after DOX induction of LZK overexpression. The changes thresholds of Log2FC ≥ |1.0| and the significance threshold of –log10 p-value ≥ 2.0 were applied to identify proteins with levels decreased (green boxed area) or increased (red boxed area) in response to the DOX treatment (A). Volcano plot displaying the log2 fold change (Log2FC, x axis) against the t-test-derived −log10 statistical (n=3) p-value (y axis) for all proteins detected in the total cell extract from CAL33 TR LZK WT cells following induction of LZK overexpression and the treatment with PROTAC97 or DMSO. The changes thresholds of Log2FC ≥ |1.0| and the significance threshold of –log10 p-value ≥ 2.0 were applied to identify proteins with levels decreased (green boxed area) or increased (red boxed area) in response to the PROTAC97 treatment (B). **C.** Venn diagram illustrating the number of overlapping upregulated proteins from DOX vs. NO DOX (yellow boxed area on volcano plot A.) and downregulated proteins from PROTAC97 *vs*. DMSO (blue boxed area on volcano plot B.) MS analysis. The proteomic analysis of protein levels is represented in data file S2.

**Figure S10.**
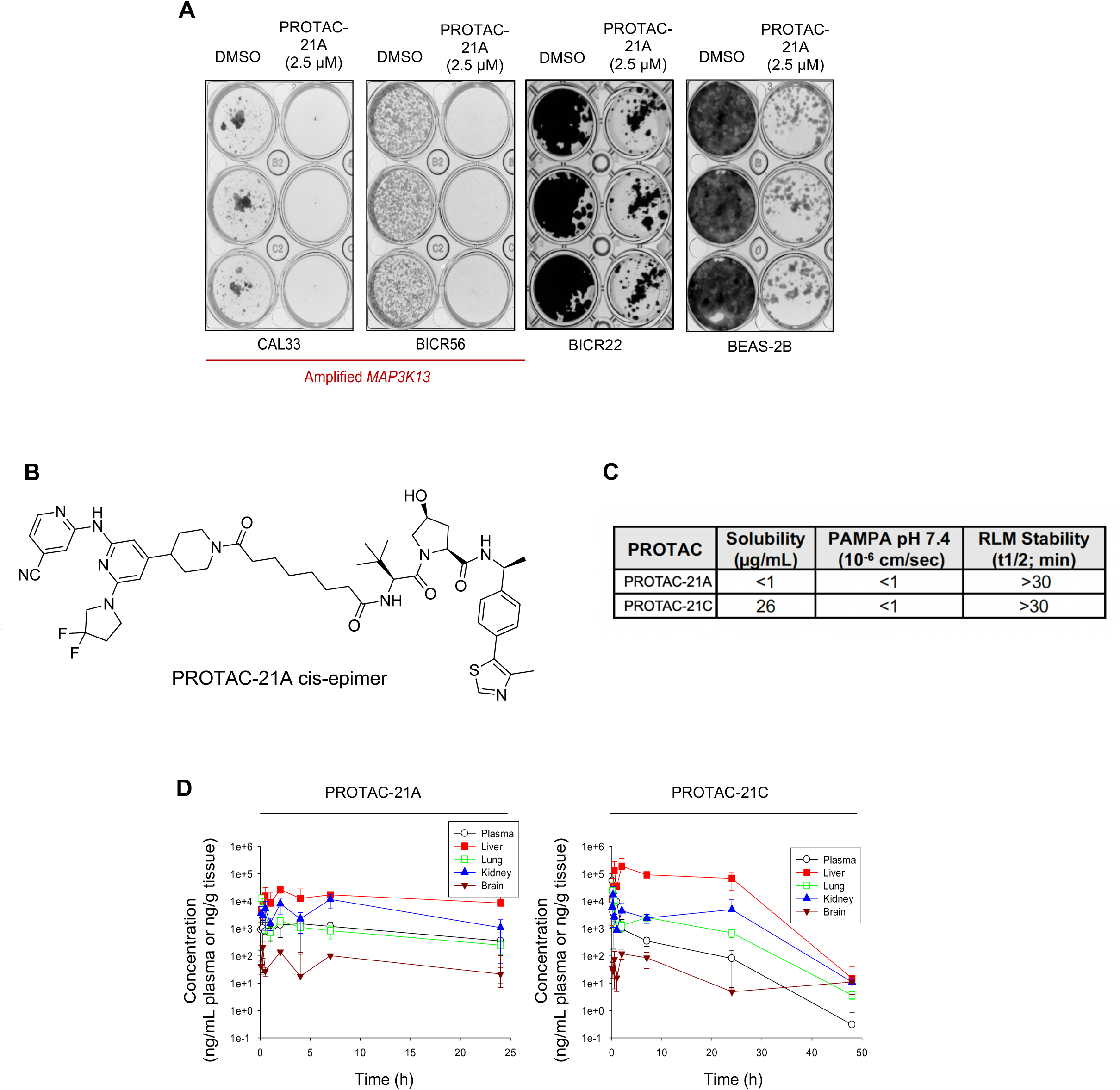
LZK-targeting PROTAC reduces HNSCC viability but exhibits low membrane permeability. **A.** Effect of PROTAC-21A on colony formation of the indicated cell lines. Data are representative of 3 independent experiments. **B.** Chemical structure of PROTAC-21A cis-epimer. **C.** *In vitro* ADME properties of PROTAC-21A and PROTAC-21C were assessed by kinetic solubility assay, parallel artificial membrane permeability assay (PAMPA) and rat liver microsomal (RLM) stability assay. **D.** Pharmacokinetic profiles of PROTAC-21A and PROTAC-21C were determined in female NSG mice after single IP administration of the PROTACs at 50 mg/kg. Mean concentration of each compound ± SD; n = 3 per time point.

**Supplementary Table S1.** *MAP3K13* amplification status for the HNSCC PDX models used in this study.

| PDX model | Institute or Company | *MAP3K13* gene expression* (FPKM) | Disease classification |
| --- | --- | --- | --- |
| 391396-364-R | NCI Patient-Derived Models Repository (PDMR) | 12.94 | Tongue (left base) |
| 295223-140-R | NCI Patient-Derived Models Repository (PDMR) | 12.43 | Neck (left) |
| 328373-195-R | NCI Patient-Derived Models Repository (PDMR) | 2.62 | Neck (left) |
| 959717-210-R | NCI Patient-Derived Models Repository (PDMR) | 1.6 | Tongue (left) |

\*High *MAP3K13* expression is defined as FPKM>5 N/A, not applicable

**Supplementary Table S2.**
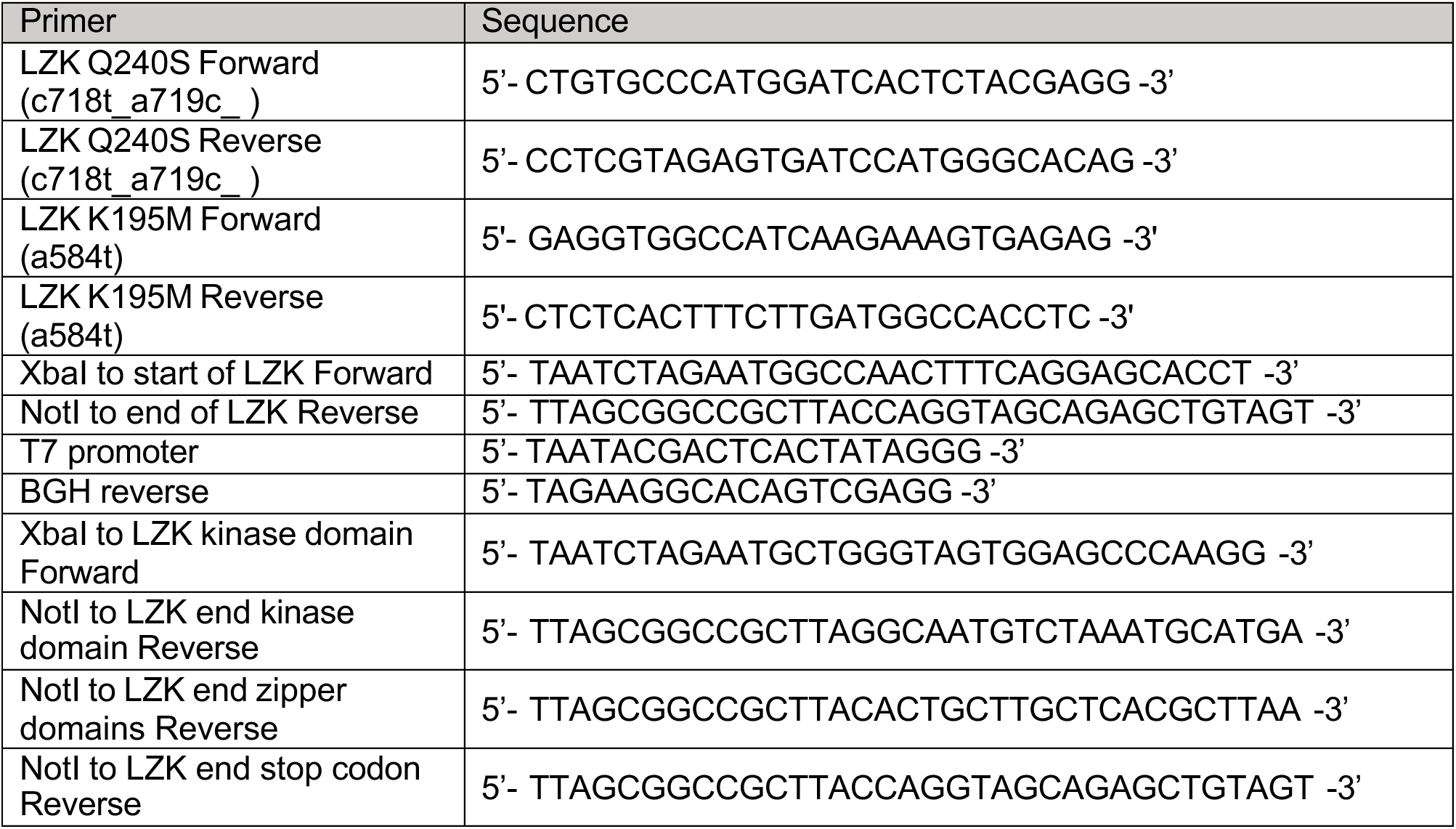
List of oligonucleotides used in this study.

**Supplementary Table S3.**
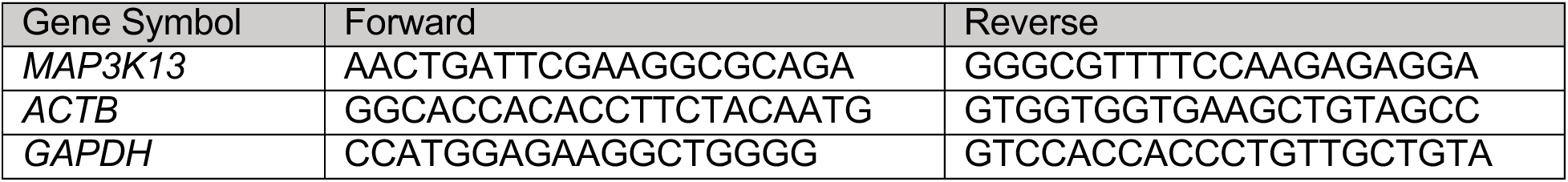
Primer sets for RT-PCR analyses.

**Supplementary Table S4.** List of antibodies used for western blot analysis.

| Antibody | Source | Identifier |
| --- | --- | --- |
| Rabbit anti-phospho-SAPK/JNK (Thr183/Tyr185) (81E11) | Cell Signaling Technology | Cat# 4668, RRID:AB_823588 |
| Rabbit anti-SAPK/JNK | Cell Signaling Technology | Cat# 9252, RRID:AB_2250373 |
| Rabbit anti-GAPDH (14C10) | Cell Signaling Technology | Cat# 2118, RRID:AB_561053 |
| Rabbit anti-phospho-MKK7 (Ser271/Thr275) | Cell Signaling Technology | Cat# 4171, RRID:AB_2250408 |
| Rabbit anti-MKK7 | Cell Signaling Technology | Cat# 4172, RRID:AB_330914 |
| Mouse anti-GST (26H1) | Cell Signaling Technology | Cat# 2624, RRID:AB_2189875 |
| Rabbit anti-c-Myc (Y69) | Abcam | Cat# ab32072, RRID:AB_731658 |
| Mouse anti-p53 (DO-1) | Santa Cruz Biotechnology | Cat# sc-126, RRID:AB_628082 |
| Rabbit anti-LZK | YenZym Antibodies | Cat# YZ6696 |
| Mouse anti-Transferrin receptor (H68.4) | Invitrogen | Cat#13_6800, RRID:AB_2533029 |
| Rabbit anti-Hsp90 (C45G5) | Cell Signaling Technology | Cat#4877, RRID:AB_2233307 |
| Mouse anti-Lamin A/C (4C11) | Cell Signaling Technology | Cat#4777, RRID:AB_10545756 |
| Mouse anti-Histone H3 (96C10) | Cell Signaling Technology | Cat#3638, RRID:AB_1642229 |
| Rabbit anti-PARP (46D11) | Cell Signaling Technology | Cat#9532, RRID:AB_659884 |
| Rabbit anti-cleaved Caspase-3 (Asp175) (5A1E) | Cell Signaling Technology | Cat#9664, RRID:AB_2070042 |
| Rat anti-FLAG (L5) | BioLegend | Cat# 637302, RRID:AB_1134268 |
| Rabbit anti-FLAG (D6W5B) | Cell Signaling Technology | Cat# 14793, RRID:AB_2572291 |
| Sheep anti-mouse IgG, secondary, HRP | GE Healthcare Life Sciences | Cat# NA931, RRID:AB_772210 |
| Donkey anti-rabbit IgG, secondary, HRP | GE Healthcare Life Sciences | Cat# NA934, RRID:AB_772206 |

## References

1. Z. C. Edwards, E. W. Trotter, P. Torres-Ayuso, P. Chapman, H. M. Wood, K. Nyswaner, J. Brognard, Survival of Head and Neck Cancer Cells Relies upon LZK Kinase-Mediated Stabilization of Mutant p53. Cancer Res 77, 4961–4972 (2017).

2. N. Cancer Genome Atlas, Comprehensive genomic characterization of head and neck squamous cell carcinomas. Nature 517, 576–582 (2015).

3. K. K. Ang, Q. Zhang, D. I. Rosenthal, P. F. Nguyen-Tan, E. J. Sherman, R. S. Weber, J. M. Galvin, J. A. Bonner, J. Harris, A. K. El-Naggar, M. L. Gillison, R. C. Jordan, A. A. Konski, W. L. Thorstad, A. Trotti, J. J. Beitler, A. S. Garden, W. J. Spanos, S. S. Yom, R. S. Axelrod, Randomized phase III trial of concurrent accelerated radiation plus cisplatin with or without cetuximab for stage III to IV head and neck carcinoma: RTOG 0522. J Clin Oncol 32, 2940–2950 (2014).

4. B. Burtness, K. J. Harrington, R. Greil, D. Soulieres, M. Tahara, G. de Castro, Jr., A. Psyrri, N. Baste, P. Neupane, A. Bratland, T. Fuereder, B. G. M. Hughes, R. Mesia, N. Ngamphaiboon, T. Rordorf, W. Z. Wan Ishak, R. L. Hong, R. Gonzalez Mendoza, A. Roy, Y. Zhang, B. Gumuscu, J. D. Cheng, F. Jin, D. Rischin, K.-. Investigators, Pembrolizumab alone or with chemotherapy versus cetuximab with chemotherapy for recurrent or metastatic squamous cell carcinoma of the head and neck (KEYNOTE-048): a randomised, open-label, phase 3 study. Lancet 394, 1915–1928 (2019).

5. M. G. McCusker, D. Orkoulas-Razis, R. Mehra, Potential of Pembrolizumab in Metastatic or Recurrent Head and Neck Cancer: Evidence to Date. Onco Targets Ther 13, 3047–3059 (2020).

6. R. Uppaluri, K. M. Campbell, A. M. Egloff, P. Zolkind, Z. L. Skidmore, B. Nussenbaum, R. C. Paniello, J. T. Rich, R. Jackson, P. Pipkorn, L. S. Michel, J. Ley, P. Oppelt, G. P. Dunn, E. K. Barnell, N. C. Spies, T. Lin, T. Li, D. T. Mulder, Y. Hanna, I. Cirlan, T. J. Pugh, T. Mudianto, R. Riley, L. Zhou, V. Y. Jo, M. D. Stachler, G. J. Hanna, J. Kass, R. Haddad, J. D. Schoenfeld, E. Gjini, A. Lako, W. Thorstad, H. A. Gay, M. Daly, S. J. Rodig, I. S. Hagemann, D. Kallogjeri, J. F. Piccirillo, R. D. Chernock, M. Griffith, O. L. Griffith, D. R. Adkins, Neoadjuvant and Adjuvant Pembrolizumab in Resectable Locally Advanced, Human Papillomavirus-Unrelated Head and Neck Cancer: A Multicenter, Phase II Trial. Clin Cancer Res 26, 5140–5152 (2020).

7. A. Cassell, J. R. Grandis, Investigational EGFR-targeted therapy in head and neck squamous cell carcinoma. Expert opinion on investigational drugs 19, 709–722 (2010).

8. J. A. Bonner, P. M. Harari, J. Giralt, N. Azarnia, D. M. Shin, R. B. Cohen, C. U. Jones, R. Sur, D. Raben, J. Jassem, R. Ove, M. S. Kies, J. Baselga, H. Youssoufian, N. Amellal, E. K. Rowinsky, K. K. Ang, Radiotherapy plus cetuximab for squamous-cell carcinoma of the head and neck. The New England journal of medicine 354, 567–578 (2006).

9. J. B. Vermorken, R. Mesia, F. Rivera, E. Remenar, A. Kawecki, S. Rottey, J. Erfan, D. Zabolotnyy, H. R. Kienzer, D. Cupissol, F. Peyrade, M. Benasso, I. Vynnychenko, D. De Raucourt, C. Bokemeyer, A. Schueler, N. Amellal, R. Hitt, Platinum-based chemotherapy plus cetuximab in head and neck cancer. N Engl J Med 359, 1116–1127 (2008).

10. J. B. Vermorken, J. Trigo, R. Hitt, P. Koralewski, E. Diaz-Rubio, F. Rolland, R. Knecht, N. Amellal, A. Schueler, J. Baselga, Open-label, uncontrolled, multicenter phase II study to evaluate the efficacy and toxicity of cetuximab as a single agent in patients with recurrent and/or metastatic squamous cell carcinoma of the head and neck who failed to respond to platinum-based therapy. Journal of clinical oncology : official journal of the American Society of Clinical Oncology 25, 2171–2177 (2007).

11. G. R. Oxnard, A. Binder, P. A. Janne, New targetable oncogenes in non-small-cell lung cancer. J Clin Oncol 31, 1097–1104 (2013).

12. J. Qian, M. Hassanein, M. D. Hoeksema, B. K. Harris, Y. Zou, H. Chen, P. Lu, R. Eisenberg, J. Wang, A. Espinosa, X. Ji, F. T. Harris, S. M. Rahman, P. P. Massion, The RNA binding protein FXR1 is a new driver in the 3q26-29 amplicon and predicts poor prognosis in human cancers. Proc Natl Acad Sci U S A 112, 3469–3474 (2015).

13. V. Justilien, M. P. Walsh, S. A. Ali, E. A. Thompson, N. R. Murray, A. P. Fields, The PRKCI and SOX2 oncogenes are coamplified and cooperate to activate Hedgehog signaling in lung squamous cell carcinoma. Cancer cell 25, 139–151 (2014).

14. P. Torres-Ayuso, E. An, K. M. Nyswaner, R. C. Bensen, D. A. Ritt, S. I. Specht, S. Das, T. Andresson, R. E. Cachau, R. J. Liang, A. L. Ries, C. M. Robinson, S. Difilippantonio, B. Gouker, L. Bassel, B. O. Karim, C. J. Miller, B. E. Turk, D. K. Morrison, J. Brognard, TNIK is a therapeutic target in Lung Squamous Cell Carcinoma and regulates FAK activation through Merlin. Cancer Discov, (2021).

15. N. Bhattacharya, A. Roy, B. Roy, S. Roychoudhury, C. K. Panda, MYC gene amplification reveals clinical association with head and neck squamous cell carcinoma in Indian patients. J Oral Pathol Med 38, 759–763 (2009).

16. B. Singh, S. K. Gogineni, P. G. Sacks, A. R. Shaha, J. P. Shah, A. Stoffel, P. H. Rao, Molecular cytogenetic characterization of head and neck squamous cell carcinoma and refinement of 3q amplification. Cancer Res 61, 4506–4513 (2001).

17. Q. Zhang, X. Li, K. Cui, C. Liu, M. Wu, E. V. Prochownik, Y. Li, The MAP3K13-TRIM25-FBXW7alpha axis affects c-Myc protein stability and tumor development. Cell Death Differ 27, 420–433 (2020).

18. S. Patel, F. Cohen, B. J. Dean, K. De La Torre, G. Deshmukh, A. A. Estrada, A. S. Ghosh, P. Gibbons, A. Gustafson, M. P. Huestis, C. E. Le Pichon, H. Lin, W. Liu, X. Liu, Y. Liu, C. Q. Ly, J. P. Lyssikatos, C. Ma, K. Scearce-Levie, Y. G. Shin, H. Solanoy, K. L. Stark, J. Wang, B. Wang, X. Zhao, J. W. Lewcock, M. Siu, Discovery of dual leucine zipper kinase (DLK, MAP3K12) inhibitors with activity in neurodegeneration models. J Med Chem 58, 401–418 (2015).

19. A. Ikeda, K. Hasegawa, M. Masaki, T. Moriguchi, E. Nishida, Y. Kozutsumi, S. Oka, T. Kawasaki, Mixed lineage kinase LZK forms a functional signaling complex with JIP-1, a scaffold protein of the c-Jun NH(2)-terminal kinase pathway. J Biochem 130, 773–781 (2001).

20. L. García-Gutiérrez, M. D. Delgado, J. León, MYC Oncogene Contributions to Release of Cell Cycle Brakes. Genes (Basel) 10, (2019).

21. S. D. Cappell, M. Chung, A. Jaimovich, S. L. Spencer, T. Meyer, Irreversible APC(Cdh1) Inactivation Underlies the Point of No Return for Cell-Cycle Entry. Cell 166, 167–180 (2016).

22. S. L. Spencer, S. D. Cappell, F. C. Tsai, K. W. Overton, C. L. Wang, T. Meyer, The proliferation-quiescence decision is controlled by a bifurcation in CDK2 activity at mitotic exit. Cell 155, 369–383 (2013).

23. E. Talevich, A. H. Shain, T. Botton, B. C. Bastian, CNVkit: Genome-Wide Copy Number Detection and Visualization from Targeted DNA Sequencing. PLoS Comput Biol 12, e1004873 (2016).

24. B. Li, C. N. Dewey, RSEM: accurate transcript quantification from RNA-Seq data with or without a reference genome. BMC Bioinformatics 12, 323 (2011).

25. R. Poplin, V. Ruano-Rubio, M. A. DePristo, T. J. Fennell, M. O. Carneiro, G. A. Van der Auwera, D. E. Kling, L. D. Gauthier, A. Levy-Moonshine, D. Roazen, K. Shakir, J. Thibault, S. Chandran, C. Whelan, M. Lek, S. Gabriel, M. J. Daly, B. Neale, D. G. MacArthur, E. Banks, Scaling accurate genetic variant discovery to tens of thousands of samples. bioRxiv, 201178 (2018).

26. I. Churcher, Protac-Induced Protein Degradation in Drug Discovery: Breaking the Rules or Just Making New Ones? J Med Chem 61, 444–452 (2018).

27. A. C. Lai, C. M. Crews, Induced protein degradation: an emerging drug discovery paradigm. Nat Rev Drug Discov 16, 101–114 (2017).

28. M. Toure, C. M. Crews, Small-Molecule PROTACS: New Approaches to Protein Degradation. Angew Chem Int Ed Engl 55, 1966–1973 (2016).

29. J. E. Burns, M. C. Baird, L. J. Clark, P. A. Burns, K. Edington, C. Chapman, R. Mitchell, G. Robertson, D. Soutar, E. K. Parkinson, Gene mutations and increased levels of p53 protein in human squamous cell carcinomas and their cell lines. Br J Cancer 67, 1274–1284 (1993).

30. M. Oren, V. Rotter, Mutant p53 gain-of-function in cancer. Cold Spring Harb Perspect Biol 2, a001107 (2010).

31. H. Solomon, N. Dinowitz, I. S. Pateras, T. Cooks, Y. Shetzer, A. Molchadsky, M. Charni, S. Rabani, G. Koifman, O. Tarcic, Z. Porat, I. Kogan-Sakin, N. Goldfinger, M. Oren, C. C. Harris, V. G. Gorgoulis, V. Rotter, Mutant p53 gain of function underlies high expression levels of colorectal cancer stem cells markers. Oncogene 37, 1669–1684 (2018).

32. P. A. Muller, K. H. Vousden, p53 mutations in cancer. Nat Cell Biol 15, 2–8 (2013).

33. K. J. Harrington, B. Burtness, R. Greil, D. Soulières, M. Tahara, G. d. Castro, A. Psyrri, I. Brana, N. Basté, P. Neupane, Å. Bratland, T. Fuereder, B. G. M. Hughes, R. Mesia, N. Ngamphaiboon, T. Rordorf, W. Z. W. Ishak, J. Lin, B. Gumuscu, R. F. Swaby, D. Rischin, Pembrolizumab With or Without Chemotherapy in Recurrent or Metastatic Head and Neck Squamous Cell Carcinoma: Updated Results of the Phase III KEYNOTE-048 Study. Journal of Clinical Oncology 41, 790–802 (2023).

34. B. Burtness, K. J. Harrington, R. Greil, D. Soulières, M. Tahara, G. de Castro, Jr., A. Psyrri, N. Basté, P. Neupane, Å. Bratland, T. Fuereder, B. G. M. Hughes, R. Mesía, N. Ngamphaiboon, T. Rordorf, W. Z. Wan Ishak, R.-L. Hong, R. González Mendoza, A. Roy, Y. Zhang, B. Gumuscu, J. D. Cheng, F. Jin, D. Rischin, G. Lerzo, M. Tatangelo, M. Varela, J. J. Zarba, M. Boyer, H. Gan, B. Gao, B. Hughes, G. Mallesara, D. Rischin, A. Taylor, M. Burian, T. Fuereder, R. Greil, C. H. Barrios, D. O. de Castro Junior, G. Castro, F. A. Franke, G. Girotto, I. P. F. Lima, U. R. Nicolau, G. D. J. Pinto, L. Santos, A.-P. Victorino, N. Chua, F. Couture, R. Gregg, A. Hansen, J. Hilton, J. McCarthy, D. Soulieres, R. Ascui, P. Gonzalez, L. Villanueva, M. Torregroza, A. Zambrano, P. Holeckova, Z. Kral, B. Melichar, J. Prausova, M. Vosmik, M. Andersen, N. Gyldenkerne, H. Jurgens, K. Putnik, P. Reinikainen, V. Gruenwald, S. Laban, G. Aravantinos, I. Boukovinas, V. Georgoulias, A. Psyrri, D. Kwong, Y. Al-Farhat, T. Csoszi, J. Erfan, G. Horvai, L. Landherr, E. Remenar, A. Ruzsa, J. Szota, S. Billan, I. Gluck, O. Gutfeld, A. Popovtzer, M. Benasso, S. Bui, V. Ferrari, L. Licitra, F. Nole, T. Fujii, Y. Fujimoto, N. Hanai, H. Hara, K. Matsumoto, K. Mitsugi, N. Monden, M. Nakayama, K. Okami, N. Oridate, K. Shiga, Y. Shimizu, M. Sugasawa, M. Tahara, M. Takahashi, S. Takahashi, K. Tanaka, T. Ueda, H. Yamaguchi, T. Yamazaki, R. Yasumatsu, T. Yokota, T. Yoshizaki, I. Kudaba, Z. Stara, W. Z. Wan Ishak, S. K. Cheah, J. Aguilar Ponce, R. Gonzalez Mendoza, C. Hernandez Hernandez, F. Medina Soto, J. Buter, A. Hoeben, S. Oosting, K. Suijkerbuijk, A. Bratland, M. Brydoey, R. Alvarez, L. Mas, P. Caguioa, J. Querol, E. E. Regala, M. B. Tamayo, E. M. Villegas, A. Kawecki, A. Karpenko, A. Klochikhin, A. Smolin, O. Zarubenkov, B. C. Goh, G. Cohen, J. du Toit, C. Jordaan, G. Landers, P. Ruff, W. Szpak, N. Tabane, I. Brana, L. Iglesias Docampo, J. Lavernia, R. Mesia, E. Abel, V. Muratidu, N. Nielsen, V. Cristina, T. Rordorf, S. Rothschild, R.-L. Hong, H.-M. Wang, M.-H. Yang, S.-P. Yeh, C.-J. Yen, N. Ngamphaiboon, N. Soparattanapaisarn, V. Sriuranpong, S. Aksoy, I. Cicin, M. Ekenel, H. Harputluoglu, O. Ozyilkan, K. Harrington, S. Agarwala, H. Ali, R. Alter, D. Anderson, J. Bruce, B. Burtness, N. Campbell, M. Conde, J. Deeken, W. Edenfield, L. Feldman, E. Gaughan, B. Goueli, B. Halmos, U. Hegde, B. Hunis, R. Jotte, A. Karnad, S. Khan, N. Laudi, D. Laux, D. Martincic, S. McCune, D. McGaughey, K. Misiukiewicz, D. Mulford, E. Nadler, P. Neupane, J. Nunnink, J. Ohr, M. O’Malley, B. Patson, D. Paul, E. Popa, S. Powell, R. Redman, V. Rella, C. Rocha Lima, A. Sivapiragasam, Y. Su, A. Sukari, S. Wong, E. Yilmaz, J. Yorio, Pembrolizumab alone or with chemotherapy versus cetuximab with chemotherapy for recurrent or metastatic squamous cell carcinoma of the head and neck (KEYNOTE-048): a randomised, open-label, phase 3 study. The Lancet 394, 1915–1928 (2019).

35. V. Girish, A. A. Lakhani, S. L. Thompson, C. M. Scaduto, L. M. Brown, R. A. Hagenson, E. L. Sausville, B. E. Mendelson, P. K. Kandikuppa, D. A. Lukow, M. L. Yuan, E. C. Stevens, S. N. Lee, K. M. Schukken, S. M. Akalu, A. Vasudevan, C. Zou, B. Salovska, W. Li, J. C. Smith, A. M. Taylor, R. A. Martienssen, Y. Liu, R. Sun, J. M. Sheltzer, Oncogene-like addiction to aneuploidy in human cancers. Science 381, eadg4521 (2023).

36. F. André, E. Ciruelos, G. Rubovszky, M. Campone, S. Loibl, S. Rugo Hope, H. Iwata, P. Conte, A. Mayer Ingrid, B. Kaufman, T. Yamashita, Y.-S. Lu, K. Inoue, M. Takahashi, Z. Pápai, A.-S. Longin, D. Mills, C. Wilke, S. Hirawat, D. Juric, Alpelisib for PIK3CA-Mutated, Hormone Receptor–Positive Advanced Breast Cancer. New England Journal of Medicine 380, 1929–1940 (2019).

37. H. Sun, P. Shah, K. Nguyen, K. R. Yu, E. Kerns, M. Kabir, Y. Wang, X. Xu, Predictive models of aqueous solubility of organic compounds built on A large dataset of high integrity. Bioorg Med Chem 27, 3110–3114 (2019).

38. H. Sun, K. Nguyen, E. Kerns, Z. Yan, K. R. Yu, P. Shah, A. Jadhav, X. Xu, Highly predictive and interpretable models for PAMPA permeability. Bioorg Med Chem 25, 1266–1276 (2017).

39. P. Shah, E. Kerns, D. T. Nguyen, R. S. Obach, A. Q. Wang, A. Zakharov, J. McKew, A. Simeonov, C. E. Hop, X. Xu, An Automated High-Throughput Metabolic Stability Assay Using an Integrated High-Resolution Accurate Mass Method and Automated Data Analysis Software. Drug Metab Dispos 44, 1653–1661 (2016).

40. P. Sriskandarajah, A. De Haven Brandon, K. MacLeod, N. O. Carragher, V. Kirkin, M. Kaiser, S. R. Whittaker, Combined targeting of MEK and the glucocorticoid receptor for the treatment of RAS-mutant multiple myeloma. BMC cancer 20, 269 (2020).

41. S. Tyanova, T. Temu, J. Cox, The MaxQuant computational platform for mass spectrometry-based shotgun proteomics. Nature Protocols 11, 2301–2319 (2016).

42. S. Tyanova, T. Temu, P. Sinitcyn, A. Carlson, M. Y. Hein, T. Geiger, M. Mann, J. Cox, The Perseus computational platform for comprehensive analysis of (prote)omics data. Nature Methods 13, 731–740 (2016).

43. M. Arora, J. Moser, H. Phadke, A. A. Basha, S. L. Spencer, Endogenous Replication Stress in Mother Cells Leads to Quiescence of Daughter Cells. Cell Rep 19, 1351–1364 (2017).

44. E. W. Deutsch, A. Csordas, Z. Sun, A. Jarnuczak, Y. Perez-Riverol, T. Ternent, D. S. Campbell, M. Bernal-Llinares, S. Okuda, S. Kawano, R. L. Moritz, J. J. Carver, M. Wang, Y. Ishihama, N. Bandeira, H. Hermjakob, J. A. Vizcaíno, The ProteomeXchange consortium in 2017: supporting the cultural change in proteomics public data deposition. Nucleic Acids Res 45, D1100–d1106 (2017).

45. Y. Perez-Riverol, J. Bai, C. Bandla, D. García-Seisdedos, S. Hewapathirana, S. Kamatchinathan, D. J. Kundu, A. Prakash, A. Frericks-Zipper, M. Eisenacher, M. Walzer, S. Wang, A. Brazma, J. A. Vizcaíno, The PRIDE database resources in 2022: a hub for mass spectrometry-based proteomics evidences. Nucleic Acids Res 50, D543–d552 (2022).

